# Graphene nanoedge electronics for monolithic and chronic recording of local microcircuits at neuronal density

**DOI:** 10.1101/2025.10.30.685707

**Authors:** Yunxia Jin, Chne-Wuen Tsai, Sippanat Achavananthadith, Dat T. Nguyen, Ze Xiong, Zhuoming Liang, Roger Herikstad, Yu Chuan, Selman A. Kurt, Hashina Parveen Anwar Ali, Benjamin C. K. Tee, Yong Lin Kong, Camilo Libedinsky, Andrew Y. Y. Tan, Chwee Teck Lim, John S. Ho, Yuxin Liu

**Affiliations:** Institute for Health Innovation and Technology, National University of Singapore, Singapore; Department of Biomedical Engineering, National University of Singapore, Singapore; The N.1 Institute for Health, National University of Singapore, Singapore; Department of Electrical and Computer Engineering, National University of Singapore, Singapore; Integrative Sciences and Engineering Programme, National University of Singapore Graduate School, Singapore; Department of Psychology, National University of Singapore, Singapore; Department of Materials Science and Engineering, National University of Singapore, Singapore; Smart Systems Institute, National University of Singapore, Singapore; Department of Mechanical Engineering, Rice University, Houston, TX 77005, USA; Department of Physiology, Yong Loo Lin School of Medicine, National University of Singapore, Singapore; Healthy Longevity Translational Research Programme, Yong Loo Lin School of Medicine, National University of Singapore, Singapore; Cardiovascular Translational Research Programme, Yong Loo Lin School of Medicine, National University of Singapore, Singapore; Neurobiology Programme, Life Sciences Institute, National University of Singapore, Singapore; Mechanobiology Institute, National University of Singapore, Singapore

## Abstract

High-density single-unit recording among closely spaced neurons over long durations is crucial for understanding the cellular-level functional architecture of the brain. Existing brain probes, however, either laterally sample neurons far more sparsely than neuronal density in the same cortical layer or suffer from long-term instability due to electrode modification, preventing precise neuron-to-neuron interrogation in local microcircuits. Here, we report a monolithic graphene-edge probe (NeuroEdge) that achieves single-unit recording at neuronal density (16 electrodes within 100 µm diameter). We fabricate NeuroEdge using self-assembled reduced graphene oxide nanoflakes to form an electrochemically active tip consisting of exposed graphene nanoedges and electrolyte-filling nanotunnels, achieving an ultralow specific impedance of 20 MΩ µm^2^. In vivo experiments over 5 months demonstrate recording at a high signal-to-noise ratio (>20 dB) and reliable interrogation of neighboring neurons. We also show that NeuroEdge can record from a single auditory cortical layer and reveal heterogeneities in the acoustic frequency response and dynamic connectivity among neighboring neurons. NeuroEdge provides a tool for precisely interrogating local microcircuits at the density of neurons in the brain.

## Introduction

Recording extracellular action potentials at high spatiotemporal resolution is crucial for understanding brain functions, investigating the pathophysiology of neurological diseases, and advancing brain-computer interfaces^1–10^. Most neuronal interactions and computations occur within spatially localized regions containing closely spaced neuron populations^11–13^. Owing to variable function, morphology, and plasticity of individual neurons, electrophysiological activity within even submillimeter-scale brain regions can be cooperative yet highly heterogeneous^14–17^. Thus, precise interrogation of complex local micro-circuits among such tightly packed neurons requires neural recording probes to reach the spatial resolution matching neuronal density, and temporal resolution matching the sub-millisecond dynamics of single-neuron activity^2,4,18^. Recent advances in silicon probe and flexible electronics technology, such as the Neuropixels probe, have enabled millisecond-resolution recording of large neuron populations^19–21^. However, the spatial resolution in the lateral direction is limited to an intershank spacing >100 µm, which is much larger than the average neuron-to-neuron distance (<25 µm) (Supplementary Fig. S1). This limitation arises from the inherent 2D planar nature of semiconductor manufacturing techniques, which restrict their ability to accurately identify single unit within cortical layer and interrogate the intralaminar connectivity of brain circuits (Supplementary Fig. S2)^3,7,19, 21–23^. To date, higher lateral density has been achieved using microelectrode arrays including Utah array^24,25^, Michigan array^26^ and large-scale microwire arrays^27,28^. However, these probes still have an interelectrode spacing ∼100 µm, 16 times sparser than neuronal density^29^. While tetrodes, some silicon probes^19,30^ or polymer composite fibers^31^ can have a few closely spaced channels, these have not been scaled to beyond 4 channels per 100 µm diameter.

A key challenge in increasing recording density is the attenuated signal-to-noise ratio (SNR) caused by the increased electrochemical impedance of electrodes as they approach cellular-scale dimensions^4,18,32–34^. Surface modification techniques are currently the only approach to address this issue and have been employed to decrease electrode impedance in both rigid silicon and flexible probes^5,7,18,22,30,35–45^. However, surface modification methods inherently limit long-term recording quality due to instability of the modified materials and their weakened bonding at electrode interfaces in physiological conditions^10,45,46^. For example, conductive polymer coatings, such as poly(3,4-ethylenedioxythiophene) polystyrene sulfonate (PEDOT:PSS) based modification, while lowering electrode impedance, are prone to delamination and degradation in chronic implantation^32,46–48^. Additionally, nanoparticle-coated platinum/iridium-based tetrodes, which are widely used for neural recording and triangulation of spike sorting, have limited effective lifetimes because of corrosion of the nanoparticle-based modification layer and associated cytotoxicity to surrounding tissue^11,49,50^. These factors limit the use of surface-modified probes for long-term, high-quality single-unit recording with minimum immunohistochemical response in many in vivo electrophysiology and neuroscience studies, where precise chronic interrogation of electrophysiological activities is essential^18,51^.

Here, we report a monolithic graphene nanoedge (NeuroEdge) probe that achieves a channel density of >16 electrodes per 0.01 mm^2^ at <25 µm interelectrode spacing, approaching cortical neuronal density, with long-term recording stability in the brain. NeuroEdge integrates more than 16 channels within a diameter of 100 µm, comparable to one human hair, using a self-assembly fabrication strategy. We achieve a neuronal size at each channel without the need for any surface modification but still maintain an ultralow impedance (20 MΩ µm^2^) which is even lower than modified electrodes. This is because we expose electrochemically active, nanotextured graphene edges and nanoporous tunnels at the probe tip. We demonstrate high-SNR (exceeding 20 dB) multichannel recording of oversampled neurons over five months, allowing high-quality spike-sorting to discriminate near-by neurons in a localized volume. We further use NeuroEdge to record from a single auditory cortical layer in mice and reveal heterogeneities of neighboring neurons, reflected by variation in preferred acoustic frequency response to sound stimulation, as well as frequency-dependent changes in functional connectivity.

### Design, fabrication and characterization of NeuroEdge

NeuroEdge array was designed to be neuronal size in each channel with equivalent circular diameter of 12-15 µm and inter-channel spacing similar to cortical neuron-to-neuron distance (Fig. 1a-c, Supplementary Fig. S1). We selected a probe diameter of ∼100 µm to eavesdrop on sufficient neurons within functional cortical columns (100-400 μm), while limiting the size of the cross-section^52,53^. As a result, the probe has 16 microelectrodes in close proximity to fill out the 100-µm-region with in the intralaminar plane, closely matching the number of neurons within the same area. The electrode density is 4-fold greater than tetrodes, 8-fold higher than Neuropixels and 16-fold higher than the Utah arrays across a 100 µm × 100 µm lateral region (Fig. 1b and f, Supplementary Fig. S1). This configuration allows each channel in NeuroEdge to access an overlapping volume of neurons, enabling triangulation-based high-quality spike-sorting of single neurons across multiple channels (Supplementary Fig. S2)^7,54,55^.

**Fig. 1.**
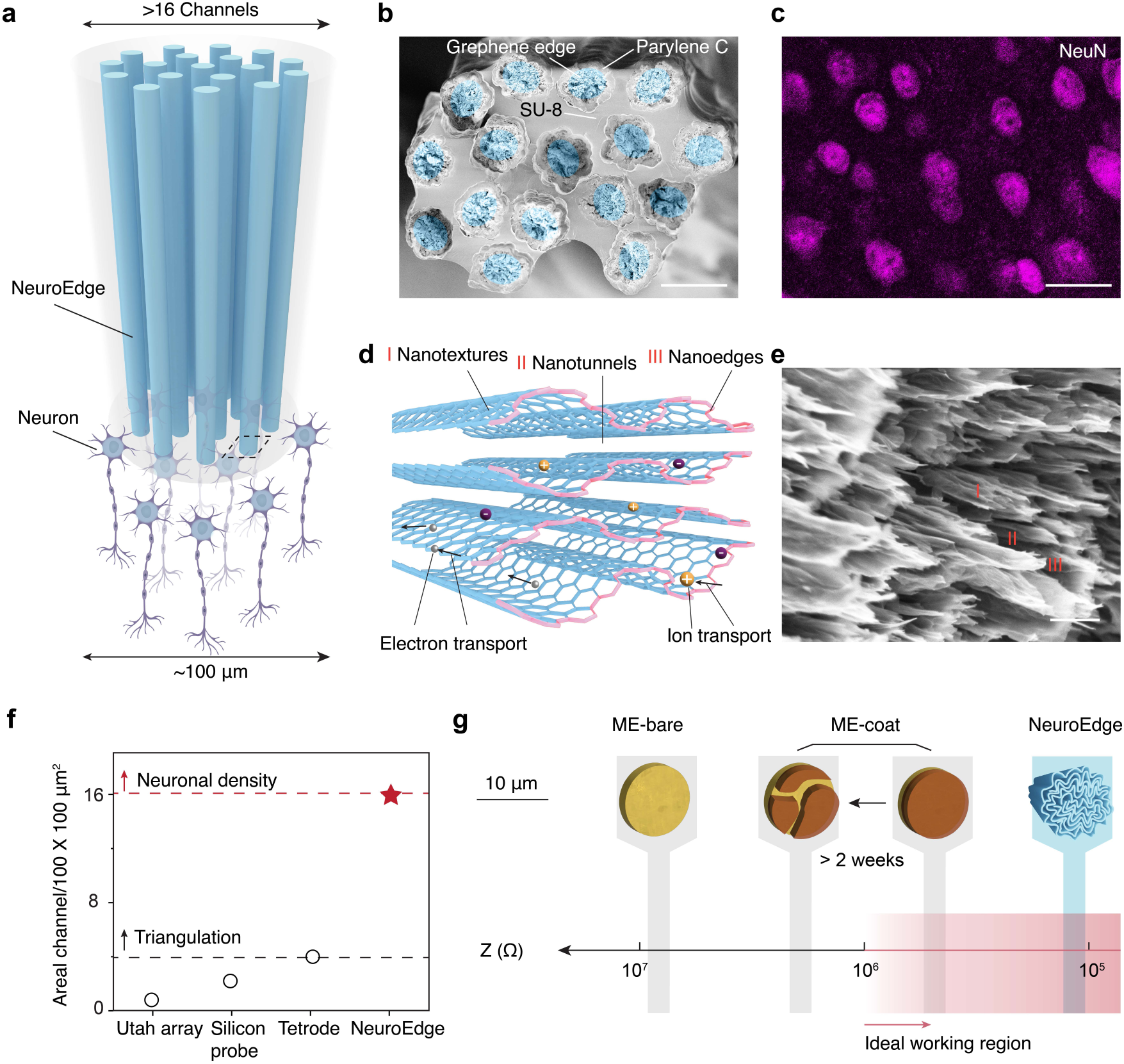
Overview of the monolithic high-density NeuroEdge. **a**, The schematic of NeuroEdge array interface with individual neurons at near 1:1 number ratio within an overall cross-section dimension of less than 100 µm. **b**, Scanning electron microscope (SEM) image of a cross-section of a 16-channel NeuroEdge array. The NeuroEdge is coated with 2-µm parylene C for insulation and the NeuroEdge array is bundled via elastocapillary force-induced self-assembly from photoresist SU-8 bath followed by UV curing. **c,** Confocal image of neuron in the cortical area of a mouse brain labelled with neuronal nuclei (NeuN). The NeuroEdge and neurons show a similar density. **d**, Schematic of NeuroEdge tip. It features nanotexture, nanoedges and nanotunnels at the tip allowing for large relative electrochemical surface area and ion/electron coupling along nanotunnels. **e**, SEM images of the cross-section of NeuroEdge. The tip of NeuroEdge is sharply exposed using a stress-concentrated breaking technique Where the I, II, III mark the nanotexture, nanotunnel and nanoedge, respectively. **f,** Comparison of the probe density of NeuroEdge, multielectrode array (MEA), and silicon probe in the lateral direction within 100 µm × 100 µm region. **g**, Comparison of impedance (Z) of NeuroEdge, metal with a coating (ME-coat) and bare metal (ME-bare) with electrode dimensions comparable to the neuron cell body (∼10 µm). The ideal impedance for electrophysiological recording is below 10^6^ Ω. NeuroEdge is within the ideal working range. The ME-coat initially falls in the working range, but moves out of the range over time due to the delamination and degradation of the modification layer. The ME-bare is far from the ideal working range. The scale bar is 25 µm in b and c, and 0.5 µm in e.

To meet our configuration design requirements, we developed a solution-processible self-assembly approach to use the nanoedge of reduced graphene oxide (rGO), instead of carbon honeycomb lattice plane, to directly interface with neurons. Specifically, in-situ 3D self-assembly of rGO with a non-toxic reductant produced a single microelectrode gel^55^, which then underwent restricted capillary shrinking driven by water-evaporation to form a straight thin microelectrode fiber. Subsequently, parylene C-coated microelectrode fibers were integrated into a bundle via facile elastocapillary self-assembly (Supplementary Figs. 3-5)^57^. These self-assembly strategies allow for high precision, versatility and biocompatibility by excluding numerous processes and toxic chemical or any organic solvent. To expose the graphene nanoedge in the cross-section of the fiber, with the aim of reducing electrochemical impedance to enable cellular-scale recording, a stress-concentrated breaking technique was developed. We constructed the stress-concentrated point by coating a water-soluble polymer, polyvinyl alcohol, on part of the rGO bundle followed by solidification in liquid nitrogen. When a rapid force is applied on the uncoated part, the bending stiffness mismatch at the coating boundary leads to a natural fracture that expose the graphene edge nanotexture with well-preserved nanoedge- nanotunnel architecture (Fig. 1d-e, Supplementary Fig. S6). In contrast, conventional exposing methods, such as scissors cutting, freezing-assisted cracking, and high-energy cutting (laser or burning) cause deformation or damage of rGO nanosheets resulting in a substantially reduced electrochemical performance or inaccurately expanded geometric size (Supplementary Fig. S6)^58,59^. The self-assemblies and stress-concentrated breaking technique provided a facile and effective approach for fabrication of ultrahigh density neural probes with arbitrary channel count.

The NeuroEdge has a significantly lower electrochemical impedance compared with bare metallic electrodes and comparable or even lower impedance compared with modified electrodes (Fig. 1g). The nanoedge-nanotunnel architecture shown in SEM images (Fig. 1e) results in high electrochemical activity relative to its exposed geometric area^60,61^. The nanoedge is electrochemically active, and nanotunnel-filling electrolyte serves as an ionic conduction pathway parallel with electronic conduction pathway which further reduce the impedance^61^. As a result, the NeuroEdge electrode, despite having a small size comparable to an individual neuron, can achieve a low impedance of < 0.2 MΩ at 1 kHz, which is 13-fold lower than platinum/iridium (Pt/Ir) electrodes, 3-fold lower than PEDOT:PSS modified electrodes on average and beyond other modified electrodes with CNT, porous TiN and IrO_2_ etc. (Fig. 1g, 2a- c)^23,31,43,47,63,64^ before recording *in vivo*. Thus, NeuroEdge has a 3.6-fold lower thermal noise compared with the Pt/Ir electrode and 1.7-fold lower compared with the Pt/Ir/PEDOT:PSS electrode.

NeuroEdge offers high electrochemical stability by eliminating surface modifications such as PEDOT:PSS coating, porous TiN deposition or platinum nanoparticle plating, which are prone to delamination or degradation under chronic electrochemical and physiological stress. As shown in Fig. 2d, NeuroEdge maintains a specific impedance of around 20 MΩ µm^2^ over 3000 min of current pulses at a charge density of 1 mC cm^-2^, which is among the typical activation thresholds for cortical microstimulation. In contrast, PEDOT:PSS coated Pt/Ir electrode exhibits a transient increase of 2.3 times in impedance within 5 min under the same current pulsing condition. This is due to the delamination of the PEDOT:PSS layer shown in the inset SEM images, where no PEDOT:PSS was left after the delivery of electrical pulses. To further assess the stability for chronic implantation purpose, we soaked the electrodes in phosphate-buffered saline (PBS) and compared the electrochemical performance over a 50-day period. Fig. 2e shows that the PEDOT:PSS coated Pt/Ir electrode exhibited over 50% impedance change and partial delamination, whereas NeuroEdge showed a consistent impedance over 50 days of soaking without obvious change.

**Fig. 2.**
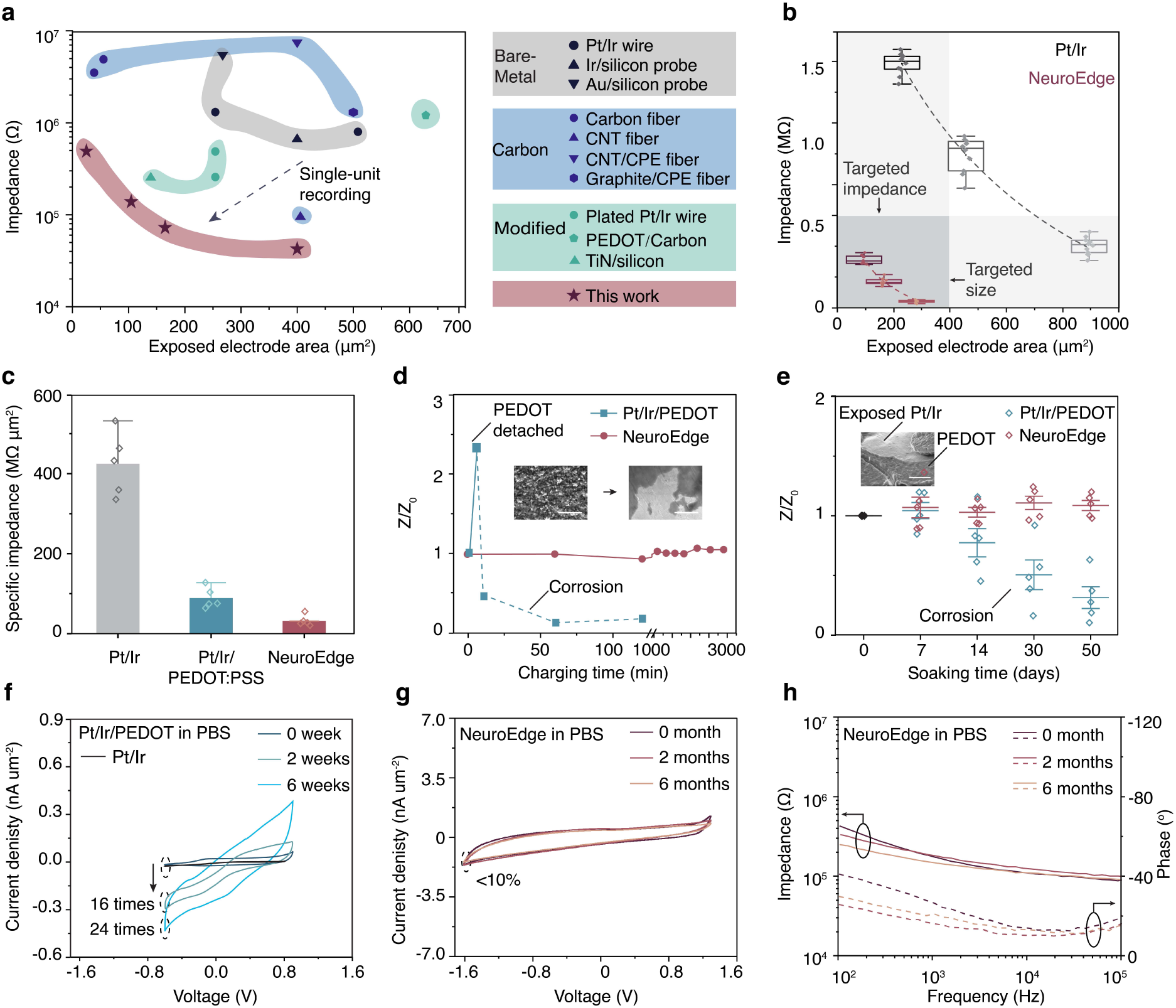
Electrochemical characterization of NeuroEdge. a, Impedance at 1 kHz of NeuroEdge compared with reported metal electrodes (Metal), surface modified metal electrodes (Modified), and carbon-based electrodes (Carbon). **b,** Impedance of NeuroEdge compared with that of conventional metal wires Pt/Ir (iridium 10%) in different diameters. **c**, Specific impedance at 1 kHz of NeuroEdge compared with Pt/Ir and PEDOT:PSS modified Pt/Ir. Values are mean ± s.d., n = 5 samples. **d**, Specific impedance at 1 kHz of NeuroEdge as a function of charging time compared with PEDOT:PSS modified Pt/Ir under a biphasic pulse. Charge density, 1 mC cm^‒2^; pulse width, 1 ms; interphase delay, 50 µs; pulse frequency, 25 Hz. **e**, Specific impedance at 1 kHz of NeuroEdge as a function of soaking time in PBS compared with PEDOT:PSS modified Pt/Ir. **f**, Cyclic voltammetry (CV) of PEDOT:PSS modified Pt/Ir at different soaking period scanning at a 100 mV S^-1^ sweep rate in the voltage range of -0.6 to 0.9 V versus Ag|AgCl. The minimum current kept dropping down by 24 times at 6-week marked in dashed circle. **g,** CV of NeuroEdge in the voltage range of -1.6 to 1.3 V versus Ag|AgCl. The minimum current was relatively stable with a change of < 10% at 6-month. **h**, Impedance of NeuroEdge over 6-month soaking in phosphate-buffered saline (PBS, 0.01 M), indicating a relatively stable electrochemical performance. The scale bar is 1 µm in d and e.

Cyclic voltammetry measurement results further confirm that the NeuroEdge has higher electrochemical stability. As seen in Fig. 2f and g, NeuroEdge has a wider water electrolysis window versus Ag|AgCl (-1.6 to 1.3 V) compared to that of Pt/Ir (-0.6 to 0.8 V). The Pt/It/PEDOT:PSS modified electrode exhibited a sharp 24-fold decrease of minimum current at the left end of the CV curve after 6-week soaking in buffer solution (Fig. 2f, Supplementary Fig. S7), while NeuroEdge only changed by 10% over 6 months (Fig. 2g). This is attributed to the PEDOT:PSS delamination and Pt/Ir corrosion. In comparison, NeuroEdge shows stable CV and EIS curves over 6 months (Fig. 2g and h). The capacitive behaviour of NeuroEdge, evidenced by the lack of redox peak in the CV curve, ensures minimal toxic side product at the electrode interface compared with redox reaction of faradaic electrodes in a physiological environment. It agrees with reported results that carbon typically has much greater resistance to electrochemical corrosion than metals used in neural probes^35^.

### High-density and high-fidelity recording *in vivo* with NeuroEdge

We assessed the recording capability of NeuroEdge by implanting it into a mouse cortex and comparing with other lateral probes (Fig. 3, Supplementary Fig. S8). After four days post- implantation, NeuroEdge registered strong electrophysiological signals. A typical 10-min raw electrophysiological data including both high-fidelity local field potential (LFP) and action potential (AP) (band-pass filtering at 300-9000 Hz) (Supplementary Figs. 9-10), remained stable over time (Supplementary Fig. S10). We also observed as high as 600 µV AP with a sharp depolarization, repolarization and relative refractory period (Supplementary Fig. S10). The power spectrum density (Supplementary Fig. S10) and the spectrogram of the recording data (Supplementary Fig. S10) provide an indication of stable recording with minimal discernible noise. Over the entire recording period, the SNR is stable as well, averaging 20 dB with maximum exceeding 30 dB (Fig. 3f). These results show that the NeuroEdge can record spontaneous action potentials with high quality.

**Fig. 3.**
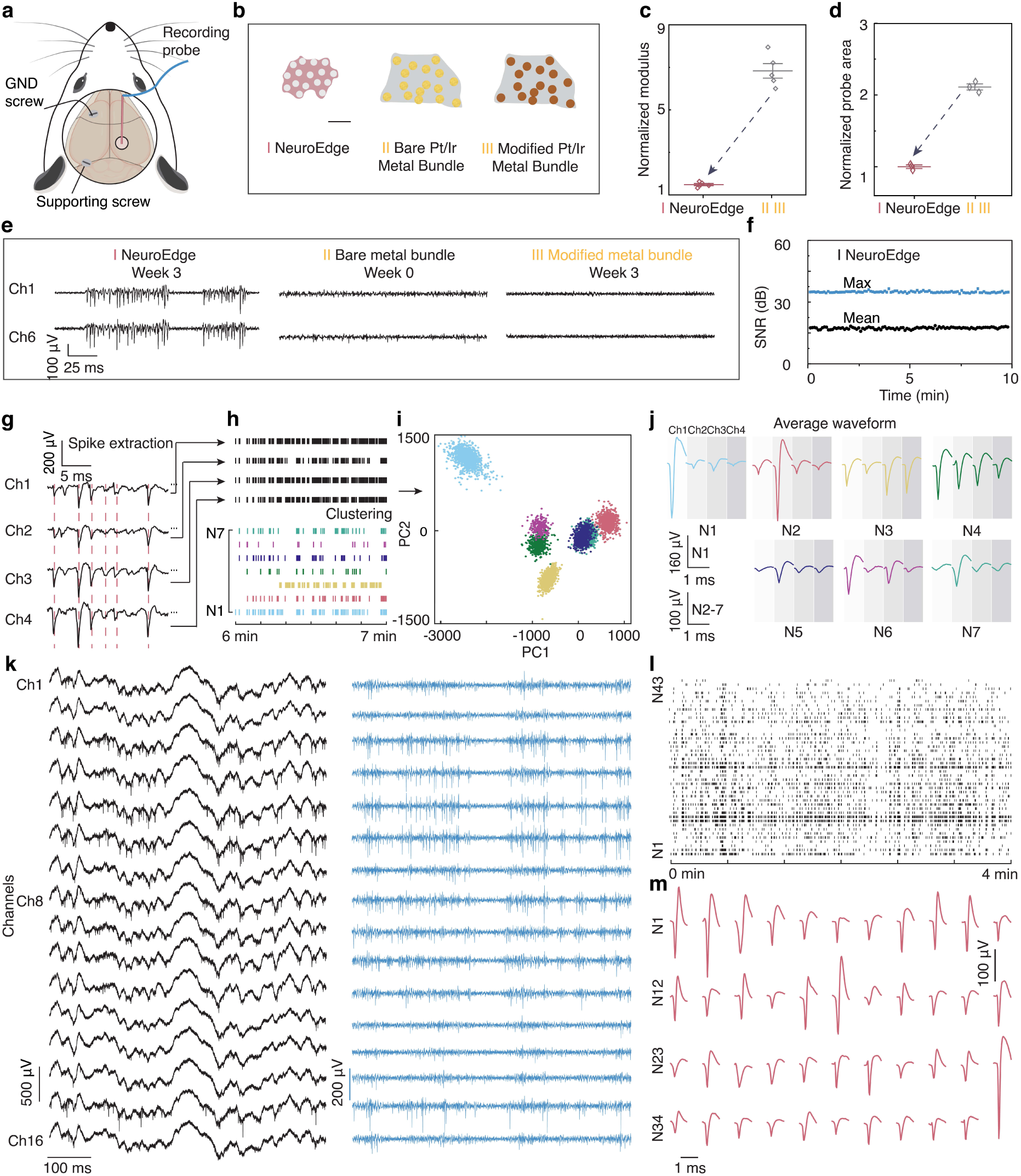
***in vivo* high-density and high-quality recording and spike-sorting with NeuroEdge. a,** Schematics of a mouse brain with implantation showing the location of 16-channel NeuroEdge, ground screw (GND), and supporting screw, as well as a 16-channel NeuroEdge neural interface (right side). **b**, Schematics of probe cross-sections. I. NeuroEdge. The white small circles represent the channel locations. II. Pt/Ir bundles assembled similarly to NeuroEdge. The gold circles indicate the channel locations. III. Microelectrode array. c, d, Comparison of these three lateral probes in terms of modulus (c) and lateral area for 16 channels (d). We set the comparison baseline to average value of NeuroEdge as one. **e,** Neural recording comparison. At week zero, bare metal wire electrode shows low SNR to allow effective recording. By week three, recordings on NeuroEdge remained stable, while almost no signal on Pt/Ir metal bundle (II) (**e**). NeuroEdge captured spikes at the same timestamp across channels. **f,** SNR in maximum and average of the recorded signal after 300-9000 Hz band-pass filtering. **g-i**, Triangulation spike sorting using Offline Sorter. It mainly includes three steps. First, filter the raw data with a second-order Butterworth HP filtering (**g**). Second, detect the waveform crossing the positive- going threshold and extract the waveform in a 1.13-ms time window with a pre-threshold period of 0.26 ms and dead time of 1.03 ms. (**h**). Spikes are stored for all four channels whenever any one of the four crosses the threshold. All the detected spikes are shown in the raster plot. Third, spike clustering based on principle components analysis (PCA) (**i**). It groups spikes with similar feature vectors that represent the activity of a single neuron. Each point on the plot represents a spike exceeding the defined threshold. Seven clusters are projected onto the first two principal components and marked as N1 to N7. **j**, Average spike waveforms on four channels for each putative neuron. **k**, Representative dense neural recording from 16-channel in cortex revealed clear and strong LFP and spiking signal (0.1-9000 Hz, left) and filtered spontaneous spiking signal (300-9000 Hz, right). **l, m,** Raster plot (**l**) and average waveforms (**m**) of the sorted units from the recording in **k**.

To compare with other lateral probes, we fabricated a Pt/Ir metal bundle (II) using the same process. The unmodified Pt/Ir bundle exhibited a lower SNR compared with that from NeuroEdge due to the high impedance, which is unable to record neural signals effectively (Fig. 3e). Due to its higher rigidity (seven times higher modulus) and larger dimensions (> two times in diameter), the PEDOT:PSS modified Pt/Ir metal lost the neural signals by week 3, while NeuroEdge remains stable recording (Fig. 3e, Supplementary Figs. 11-12). Owing to the high spatial density and high SNR, NeuroEdge is capable of oversampling the same set of neurons across more channels. In particular, it resolves both strong nearby spikes and low-amplitude spikes that would otherwise be obscured by insufficient lateral resolution or thermal noise associated with high impedance.

Four representative channels were selected to demonstrate the spike sorting process. (Fig. 3g and h). Individual neurons are simultaneously recorded across all four channels, as evidenced by similar cross-channel firing patterns with similar timestamps, which provides the basis for high-quality triangulation. The spike clustering projections are illustrated in Fig. 3i and Supplementary Fig. S13, showing 2.25 ± 0.75 single units per channel on average. Principal components analysis (PCA) reveals seven clearly separated clusters in three projections after removing sparsely firing units, multi-units, and noise units. The clusters that overlap in the PC1/PC2 projection become completely distinguishable when observed in either the PC1/PC3 or PC2/PC3 projections (Supplementary Fig. S13). The average waveforms of each unit on all four channels exhibit similar spike shapes yet varying amplitudes, suggesting that the spikes were from the same neuron but at different distances to different channels (Fig. 3j). The interspike intervals (ISI) histogram of all seven units feature a refractory period of more than 2 ms that render them consistent with single-unit activities.

Following similar spike sorting process, single units were discriminated from a 16-channel recording (Supplementary Figs. 14-15). All 16 channels captured comparable LFP with similar wave patterns, and the AP with closely aligned spike timings (Fig. 3k), indicating a highly localized high-density electrophysiological recording. The identified single units had no identical overlap in spike timings (Fig. 3l) and showed distinct average waveforms (Fig. 3m), suggesting an absence of duplicated units. As a result, this 100-µm array successfully identified over 40 well-isolated single units, enabling a laminar recording at a density exceeding 4000 units/mm^2^, approaching the neuronal density^64^.

### Investigating heterogeneity of frequency preference among neighboring neurons in auditory cortex

Frequency is the main acoustic parameter that is thought to be topographically mapped in most of the auditory cortex. Studies have identified fine-scale frequency architectonic inhomogeneity across cortical laminae in the auditory cortex^66,67^. However, the existence of local heterogeneity in frequency preference across neighboring neurons within a cortical layer remains inconclusive or incomplete^68^. This is largely because there is a lack of tools or methods that combine both high temporal resolution and high lateral spatial resolution (matching cellular density). To address this challenge, we implanted NeuroEdge (with sub-25 mm spatial resolution) into mouse auditory cortex to record closely spaced neurons in response to tone frequencies spanning 0.5-42 kHz in ½ octaves (Fig. 4a, Supplementary Fig. S16). The recorded local field potential (LFP) shows an increased negative deflection near tone onset, which is sustained for the entire 100-ms stimulus window (Supplementary Fig. S17). The filtered spiking signal reveals that all the neurons have unequal responsiveness to different sound frequencies with firing rate ranging from 0-16 Hz (Fig. 4b-c).

**Fig. 4.**
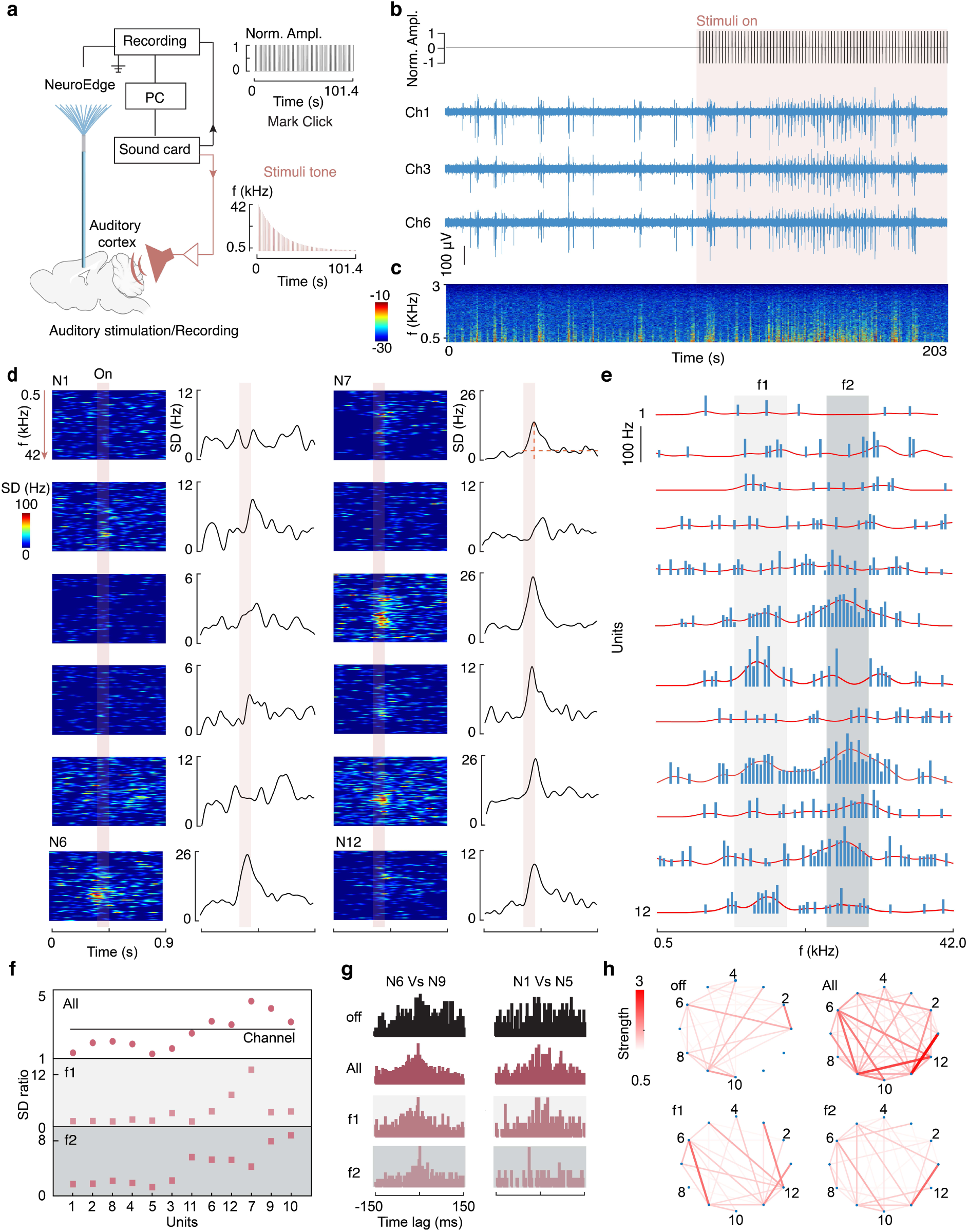
NeuroEdge resolves heterogeneity in the acoustic frequency response of neighboring neurons in mice auditory cortex. a, Schematic set-up for the auditory stimuli and recordings. The sound card supplies two channels. One is the click channel to align the timeline of the electrophysiological recording and the stimuli, and another is the tone channel to deliver auditory stimuli. There are 78 frequencies spanning 0.5-42 kHz in ½ octave with a duration of 100 ms and interstimulus interval of 1.3 s. **b, c,** Representative electrophysiological recording (**b**) and the spectrogram (**c**) from NeuroEdge during pre-stimulation and stimulation spanning 0.5-42 kHz. **d,** Time-domain spike density (SD) colormaps and average spike density curves of representative putative neurons in a time window crossing the stimuli duration. **e**, Frequency-domain stimuli response of the same representative neurons in **d**. The red lines are the fitted curves. **f,** Average SD ratio for each putative neuron responding to full frequency from 0.5 to 42 kHz (All), lower frequency marked in f1 and higher frequency marked in f2 in **d**. The center frequency of f1 and f2 is 2.6 kHz and 9.5 kHz, respectively. The black solid line represents the SD ratio from the whole channel including all the neurons. **g,** Representative cross- correlograms of neurons under no stimulation and stimulation with full frequency of 0.5-42 kHz, f1and f2 **h,** Cross-correlation matrices of pairwise neurons under no stimulation and stimulation with full frequency of 0.5-42 kHz, f1 and f2.

Fig. 4d and Supplementary Fig. S18 illustrate the characteristic frequency response in peristimulus raster plot from 12 representative sorted single units from 4 channels. It shows an average 2.7-fold increase in spike density, computed as the ratio of peak over baseline, during the stimulus window. However, all units exhibit different spike density increases ranging from 1 to 5 (Fig. 4d-f). For instance, unit 7 (N7) shows a 4.4-fold spike density increase, while unit 1 (N1) exhibits almost no change. The observed heterogeneity in spike density changes among spatially adjacent neurons within the stimulus window indicates that those nearby neurons are not uniformly activated by the external acoustic stimulus.

By observing neural responses to acoustic stimuli at varying frequencies, we found that all the units share common best frequency peaks around 2.6 kHz and 9.5 kHz, indicating they are from a vicinity region (Fig. 4e, Supplementary Fig. S19). However, each unit has its own frequency preference. For instance, units 7 and 12 (N7, N12) show strong responses to low frequencies (<4 kHz), while units 6, 10 and 11 (N6, N10, N11) show elevated responses to higher frequencies (>4 kHz), and unit 9 (N9) presents responses both below and above 4 kHz (Fig. 4g). Such frequency preference suggests heterogeneous responsivity among nearby neurons, agreeing with previously reported results using optical methods^22^. We quantify the responsivity difference under different frequency ranges by calculating the SD ratio of neurons under acoustic stimuli at lower frequency f1 centering 2.6 kHz and higher frequency f2 centering 9.5 kHz (Fig. 4f). We found a distinct preference of the units to f1 and f2. Particularly, N7 showed preference to f1 (lower frequency) with an SD ratio of 13.2 at f1, which is three times that at f2 of 4.2. In contrast, N10 showed opposite preference at f2 (higher frequency) with an SD ratio of 8.7 compared to 3.2 at f1.

We further investigate whether and how functional connectivity of nearby neurons in single cortical layer is modulated by varying acoustic frequencies. From the examination of cross-correlograms between N6 and N9 (Fig. 4g and Supplementary Fig. S20)^8^, a discernible peak at -3 ms was observed under full frequency (0.5-42 kHz) stimuli, suggesting increased excitatory functional connectivity due to the acoustic stimulation. This augmented excitatory connectivity persists at both f1 and f2 frequency bands. However, pairwise analysis of N1 and N5 reveals frequency-specific connectivity changes, notably an excitatory functional connection at the f1 frequency band and an absent connection at the f2 frequency band. Examining the interaction strength (coefficient ratio) reveals a marked increase in both the number and strength of functional connections under acoustic stimulation (Fig. 4h). Notably, interaction strength exhibits significant variability when comparing lower versus higher frequency ranges.

These results demonstrate the capability of NeuroEdge probes to discriminate substantial heterogeneity of neuronal tuning at the cellular level within a localized cortical region which has been previously understudied, and reveal the dynamic functional connectivity with frequency specificity among neighboring neurons within a functional microcircuit in the auditory cortex. NeuroEdge provides a versatile tool for investigating the neural basis of behavior, extending beyond organization across the depth of the cortex to within the intralaminar cortical layer.

### Chronic *in vivo* biocompatibility and functionality assessment

We investigated the chronic functionality of NeuroEdge by evaluating the stability of recorded electrophysiological signals, the acoustic frequency response of single units, and the biocompatibility of the implanted probe in the mouse brain (Fig. 5a). Representative electrophysiological recordings at month 1 (M1), month 3 (M3), and month 5 (M5) all exhibit prominent and high-amplitude action potential. (Fig. 5b). The SNR was consistently above 20 dB with a noise level below 7 µV across the entire experimental duration of 6 months (Fig. 5c). The average spike amplitude remained stable. NeuroEdge with size comparable to neuron also maintained a low impedance *in vivo* of around 0.5 MΩ over 6 months (Fig. 5d). The sustained high SNR and low impedance of NeuroEdge enable neural activities monitoring with high- fidelity over an extended period without deterioration in yield and quality.

**Fig. 5.**
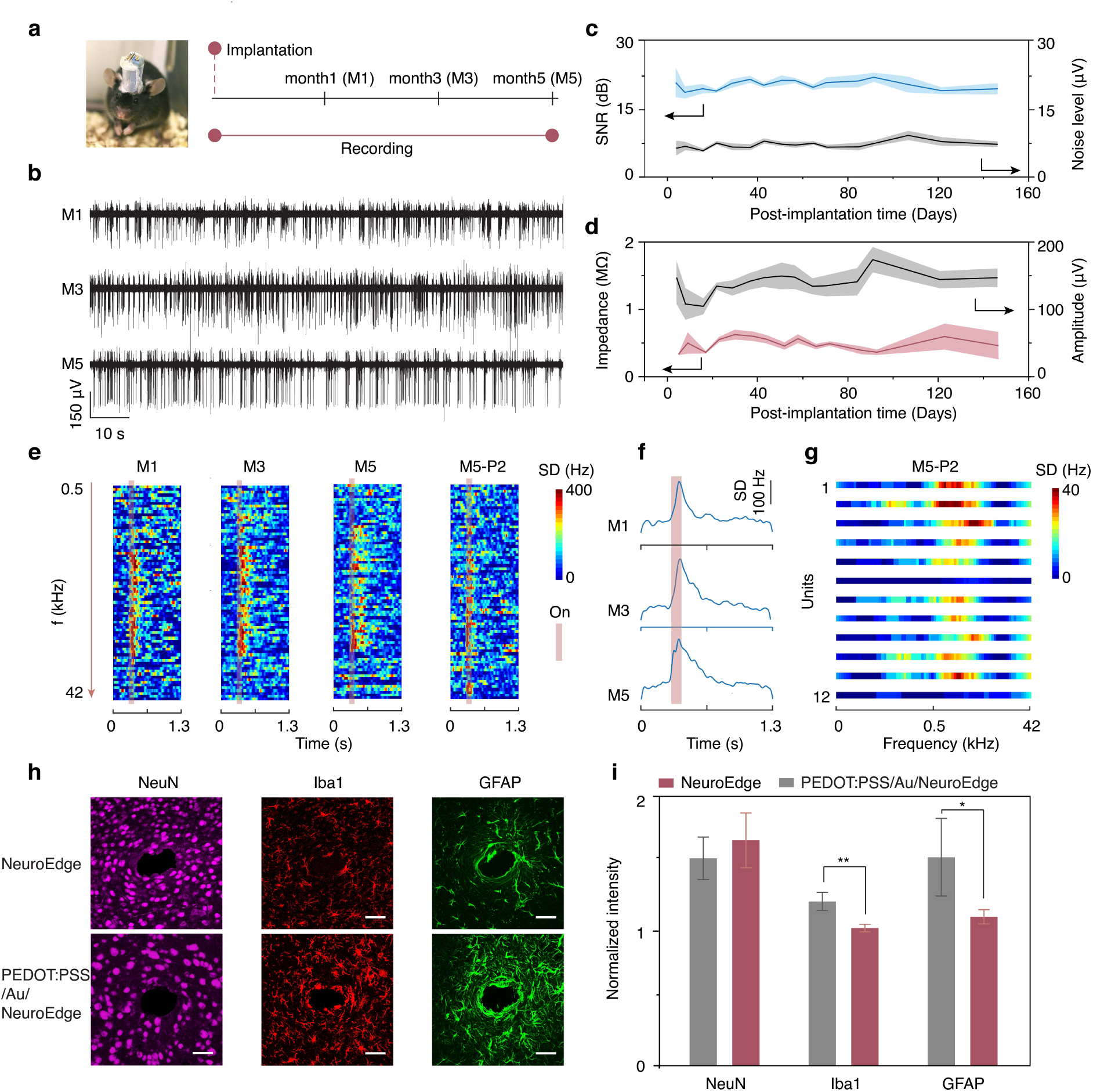
Chronic assessment of recording stability, functionality and biocompatibility of NeuroEdge. a, A photograph of a mouse implanted with NeuroEdge and a schematic of the chronic recording timeline. **b,** Representative electrophysiological recording from NeuroEdge at M1, M3 and M6. BP filtering of 300-9000 Hz was applied. **c-d**, Performance of the electrophysiological recording stability. The average values of SNR and noise level (**c**), NeuroEdge impedance and spike amplitude (**d**) all remained stable for 6 months. **e**, Peristimulus spike raster plot (**e**) and and month 5 after moving down the probe by ∼150 µm. **f,** Spike density in response to 78 frequencies spanning 0.5-42 kHz at month 1, 3, 5. **g**, Frequency response distribution of representative near-by single units in mice at M6. **h**, Chronic- biocompatibility of pristine NeuroEdge and PEDOT:PSS/Au coated NeuroEdge after 3-month implantation. Neurons, microglia and astrocytes are immunohistochemically stained with NeuN (red), Iba1(red) and GFAP (green), respectively. **i**, Analysis of the neuron cells (NeuN), activated microglia (Iba1) and astrocyte cells (GFAP) around insertion position across the entire immunohistochemical images. Values are shown mean ± s.d. using one-way t-test (*P<0.05, **P<0.01). n= 6 samples.

We further investigated the chronic functionality of NeuroEdge in response to acoustic frequency. The frequency preference of neurons has negligible change over five months, and the spike density increase remains unchanged (∼ 2.6-fold increase for M1, M3 and M5) (Fig. 5e and f). In addition, NeuroEdge remains functional and can resolve frequency preference heterogeneities in the auditory cortex, 5-month post-implantation (Fig. 5g). Altogether, the results demonstrated the stable functionality of NeuroEdge for recording neural activity and successfully discriminating diverse response of neurons to auditory stimuli over 6 months.

To access long-term biocompatibility, we compared the neuroinflammatory responses to pristine and modified neural interfaces in mouse cortex, specifically using monolithic NeuroEdge and PEDOT:PSS/Au-modified probe of similar size. Immunohistochemical studies were performed on horizontal brain slices near the implantation sites to assess the presence of neuronal nuclei, microglia cells and astrocyte cells via immunostaining with neuronal nuclei (NeuN), ionized calcium-binding adaptor molecule 1 (Iba1, positive marker for microglia cell) and glial fibrillary acidic protein (GFAP, positive marker for astrocyte cell) (Fig. 5h). We observed no significant morphology change of NeuN and Iba1 stained cells in the area surrounding implanted NeuroEdge sites at 3-month post-implantation, and the intensity values are in the same range as the non-implanted contralateral cortex section (Fig. 5i). The good biocompatibility of NeuroEdge can be ascribe to the chemical inertness and small tissue lateral displacement (<100 µm). However, we observed a 20 % increase in Iba1 intensity and 40% in GFAP intensity in the area surrounding the PEDOT:PSS/Au coated probe compared to the monolithic NeuroEdge, suggesting glial scar encapsulation of microglia cells and astrocyte cells that were triggered by the modified implant. This could be due to PEDOT:PSS degradation/disintegration in physiological solutions and poor interface adhesion between metal and PEDOT:PSS, resulting in delamination or diffusion of disintegrated dopants and unreacted monomers^32^. Overall, the chronic immunostaining study indicates that the NeuroEdge is suitable for long-term implantation without substantial neuroinflammatory response.

## Conclusion and outlook

We have reported NeuroEdge, a monolithic graphene nanoedge neural probe, that provides a recording resolution laterally matching brain neuronal density. In contrast with existing multielectrode technologies, such as the Utah array and Neuropixels, which sparsely sample neurons in the intralaminar direction, NeuroEdge achieves an ultrahigh electrode density at > 16 electrode within a region 100 × 100 µm^2^, enabling high-quality discrimination of closely packed neighboring neurons. Unlike state-of-the-art silicon probes that rely on planar processing steps, NeuroEdge is monolithically self-assembled. The monolithic design uses active graphene nanoedges and capillary nanotunnels to interface neuron, achieving a low impedance at neuronal size without any surface modification. It enables long-term and high-quality single-unit recordings (SNR > 20 dB) by eliminating the corrosion or delamination of modified layers. In mouse auditory experiments, NeuroEdge resolves heterogeneities among neighboring neurons by identifying variation of preferred acoustic frequency response and functional connectivity of each neuron. These results demonstrate the utility of NeuroEdge as a tool to precisely interrogate intralaminar microcircuitry and unravel the cellular architecture function of the brain.

Beyond the auditory cortex, we envision that NeuroEdge, which has the capability of millisecond temporal resolution, cellular-scale spatial resolution, and chronic stability, will facilitate the understanding of neural microcircuits and their functional architecture in many brain regions, especially those with dense neuron nuclei or fine-grained functional division, such as the hippocampus, visual cortex and thalamus^70^. It can be broadly applied to investigate functional connectivity and plasticity in microcircuits and how they are affected by neurological disorders by tracking many individual neurons in a localized area. While one NeuroEdge bundle contains 16 channels, multiple bundles can be implanted to expand the total channel count and investigate the functional interconnection among neighboring neurons at multiple brain regions simultaneously. The capabilities of NeuroEdge also position it as a highly promising solution for clinical bioelectronics applications. Particularly, it could facilitate the development of brain-computer interfaces, where precise control of closely spaced neurons is required for the restoration of fine motor and sensory functions^71^. Additionally, compared to existing silicon or metal-based probes, NeuroEdge can be further developed into a multifunctional neural probe, with simultaneous recording of electrophysiological and neurochemical signals, as well as localized neuromodulation by leveraging the inherently exceptional electrochemical activity of graphene edges.

## Methods

### Fabrication of graphene edge probes (NeuroEdge)

Aqueous dispersion of graphene oxide (GO) (Suzhou Tanfeng Graphene Technology CO., LTD, 10 mg ml^-1^) was diluted to 6 mg ml^-1^ with water. Then, 85 mg of ascorbic acid (Sigma-Aldrich) was added to 1 ml of the GO dispersion and ultrasonicated for 5 min. The resulting GO/ascorbic acid mixture dispersion was loaded into glass capillary tubing, sealed with Teflon tape, and reacted at 95 °C for 1 h, resulting in a reduced graphene oxide (rGO) gel fiber in the capillary tubing. The gel fiber was extracted, and soaked in a water bath for 10 min, followed by changing the water and repeating for another 10 min. The gel fiber was then placed on a flat plastic polystyrene substrate to constrict the self-assembly alignment of rGO fiber during water-evaporation driven drying. The diameter of the rGO fiber was determined by the inner diameter of the glass capillary tubing, with a diameter of 10-15 µm synthesized from 200 µm of capillary tubing. To facilitate the drive assembly, a 50 µm stainless steel wire (793400, A-M systems) was connected to the fiber using silver nanowire ink. The bare rGO fiber was coated with a 1-2 µm thick of parylene C to insulate the side wall of the fiber. The tip of the fiber was exposed via a strain- concentrated breaking. First, optimal cutting temperature (OCT, PVA solution) compound was coated on the surface of parylene C-coated rGO fiber, and frozen in liquid nitrogen. The OCT liquid solidified at the low temperature, encapsulating the entire rGO fiber, leaving a short fiber exposed at the tip end. Second, using spring scissors to press down on the free tip end, the fiber was broken down from the point adjacent to the OCT embedding part. The fiber was taken out from liquid nitrogen and the melted liquid OCT was then washed away with water. After exposing the tip end of the rGO fiber, the sample was referred to as a graphene edge probe (NeuroEdge).

### Fabrication of NeuroEdge array

The rGO fibers coated with parylene C were bundled with elastocapillary self-assembly by withdrawing the free-moving end of the fibers from the bath of diluted SU-8 solution (SU-8 2002, Kayaku Advanced Materials, Inc.). The assembly was baked in an oven at 100 °C for 10 min to evaporate the solvent in SU-8 and then exposed to a 365 nm UV lamp for 5 min to bond the rGO fibers permanently to a thin array. The tip end of the array was exposed using the strain-concentrated breaking method as described in the previous section.

### Assembly of implantation micro-drives

The stainless steel wire that is connected to NeuroEdge was tightened inside the receptacle (ED85100-ND, Digi-key) by putting back the original pin. Two holes in the receptacle were left to connect with the reference and ground wires for in vivo electrophysiological recording. The moving stage inside the trimmer potentiometer was exposed to place the PI tubing part of the NeuroEdge array which was secured with adhesive (Loctite 404, Ted Pella). To create a strong bond of NeuroEdge on the moving stage, the adhesive was left overnight to allow full curing. The NeuroEdge tip end was adjusted to pass the bottom of the potentiometer by 1-2 mm to facilitate implantation into the brain. The receptacle was secured on the top of the trimmer potentiometer. Moreover, the stainless steel wires were left bent to facilitate the moving down of the NeuroEdge.

### Characterize the exposed tip area of NeuroEdge

The size and morphology of the NeuroEdge tip end were analyzed with a Field-emission Scanning Electron Microscopy (FESEM) (FEI Verios460). The NeuroEdge array was attached to a 70° SEM stub with its tip slightly higher than the top side and grounded using silver ink on the bottom end of NeuroEdge to ensure electrical connection to double-sided carbon conductive tape adhered on the stub. The sample was not coated with additional Pt or Au to allow detection of the nanoscale graphene edges, the nanotextures and the nanotunnels. The sample holder stage was tilted to allow for the tip end to be viewed vertically or at a 45-degree angle relative to the FESEM detector.

### Mechanical characterization

Tensile testing experiments were performed using an Instron instrument (Instron model 68SC- 1). Stress-strain curves were obtained under ambient conditions with 10-mm-long samples at a constant strain rate of 1mm/min. Values of Young’s modulus were determined based on data from at least five samples. Bending stiffness (D) of the probe is defined as

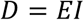

where *E* and *I* represent the elastic modulus and moment of inertia, respectively. For a cross-sectional area A, the moment of inertia is calculated as

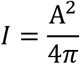

### Characterization of NeuroEdge in PBS solution

The electrochemical performance of NeuroEdge was characterized in a phosphate buffer solution (PBS, 150 mM) using an Ag/AgCl electrode as a reference and a platinum mesh as a counter. Electrochemical impedance spectroscopy (EIS) was conducted over a frequency range of 100 to 100k Hz with a perturbation of 10 mV RMS sinusoidal excitation voltage (Biologic SP200). Cyclic voltammetry (CV) curves were acquired at a sweep rate of 100 mV/s between negative and positive potential limits of -1.6 and 1.3 V versus Ag/AgCl. The current pulse was delivered by applying a rectangular charge-balanced biphasic current pulse with the cathodic phase applied first which is a common practice for in vivo stimulation (AM4100, A-M Systems). The interpulse delay between the cathodal and anodal phase was set to 50 µs to visualize the interpulse potential. The pulse width was 1 ms, and the charge density was controlled to be 1 mC cm^-2^. The long-term stability study of NeuroEdge was performed by soaking it in 150 mM PBS for half a year.

### PEDOT:PSS modified metal electrodes as control group

A metal electrode was electroplated with poly(3,4-ethylenedioxythiophene): poly(styrene sulfonate) (PEDOT:PSS) from a 0.1 M PSSNa (Sigma-Aldrich) aqueous solution with 0.01 M EDOT (Sigma-Aldrich) under galvanostatic conditions using an Ag/AgCl electrode as a reference and a platinum wire as a counter. The current density was controlled to be around 0.4 mA/cm^2^ to allow slow electrodeposition for enhanced adhesion and total electrodeposition charge density was around 600 mC/cm^2^ to minimize the risk of PEDOT:PSS layer cracking. After electrodeposition, the electrode was immersed in DI water for 15 min to wash away any weakly attached PEDOT:PSS on the surface. The electrochemical performance was characterized using EIS, CV and CIC measurements, with potential limits of -0.6 to 0.8 V versus Ag|AgCl, following the same process as the NeuroEdge characterization.

### Ethical approval and animal handling

All animal procedures are approved by the Institutional Animal Care and Use Committee (IACUC) at National University of Singapore and were conducted in accordance with NACLAR guidelines. Male C57BL/6 wild-type mice aged 8-12 weeks were purchased from InVivos, and housed in a room maintained for a 12-hour light/dark cycle with ad libitum access to food and water. Before surgery, the mice were group-housed with 4 mice per cage while after surgery, the mice were individually housed with toys and brown paper provided in their cages. The mice were allowed to acclimate for at least 3 days before being used in the experiments.

### Chronic implantation of NeuroEdge array to parietal lobe cortex

Mice were anesthetized via intraperitoneal injection of ketamine/xylazine mixture in saline (100 and 10 mg per kg mouse, respectively), and subcutaneous injection of analgesic buprenorphine (0.1 mg per kg mouse). The depth of anesthesia was monitored with the toe pinch reflex. The top of the head of the mice was shaved, and then placed in a stereotaxic frame (David Kopf Instruments), with ear bars securing the head and gel applied on the eyes to keep them moist. The exposed skin on the top of the head was cleaned with alcohol and iodine, and cut along the midline to expose the skull. After cleaning the skull with saline, two small holes were drilled, one on the right forehead and one on left part of back head for metal supporting screws (3SSCS94F008, ¼-28, Fastenere) using 0.9 mm diameter burr (19007-09, Fine Science Tools) with microdrill (78001, World Precision Instruments), respectively. Before mounting the forehead screw, a stainless-steel wire was soldered to the screw head for grounding and reference that would be connected to the electrode drive later. A craniotomy was performed on the right skull for electrode implantation using a 1.8 mm diameter trephine (18004-18, Fine Science Tools). The prepared drive with NeuroEdge was secured on an insertion tool that was mounted on the stereotaxic frame holder. The drive was then lowered down to the brain with the NeuroEdge tip facing towards the coordinates relative to bregma of -1.9 mm AP, 1.4 mm ML,

0.5-1.0 mm DV. The dura was partially removed surrounding the NeuroEdge penetration point, and sterile saline was applied to keep the craniotomy moist. The drive was fixed onto the skull using cement (C&B Metabond, Parkell Inc.) with Kwik-Sil (World Precision Instruments Inc.) applied to the craniotomy area. After the cement and Kwik-Sil had completely cured, the insertion tool was removed, and a plastic tubing was placed outside the drive to protect it. The drive and plastic tubing were then secured together on the skull with dental cement (Rapid repair Meliodent), and the gap between them on the top was sealed as well to keep the craniotomy isolated from the ambient environment. The mice with implantation were returned to their cage after waking up from anesthesia. The mice were closely monitored for 3 days after surgery.

### In vivo electrophysiological recording

An Intan recording system, which includes an RHD USB interface board and RHD headstage with an RHD2132 chip (Intan Technologies), was connected to the receptacle connector of the drive using an in-house adaptor. The mice were maintained at an anaesthetized state using 1-2 % isoflurane (Baxter Healthcare Corporation). This helped to minimize the trial-to-trial variation arising from the effects of attention, arousal, and body movements to facilitate the performance investigation of electrode itself. To minimize the noise from the environment, the mice were placed inside a Faraday cage, and the cage was grounded via the ground connector in the USB interface board. The RHX Data Acquisition Software was used to record the local field potential (LFP) and action potential (AP). The data acquisition was performed in the frequency width of 0.1-9000 Hz with a sampling rate of 30 kHz, and the in vivo impedance was measured at 1 kHz before recording.

### Auditory stimulation and recording

For auditory stimulation recording, the NeuroEdge was implanted into the left-side auditory cortex (AC) at coordinates relative to bregma of -2.2 mm AP, -4.0 mm ML and 0.6 mm DV, using the same recording and implantation setup as previously described. Pure tones were generated with a sound card (Sound Blaster Audigy Rx) and MATLAB software, and amplified via an audio amplifier (Samson Servo 120a) before being presented to the mice’s ear via loudspeaker (XT25TG30-04, Peerless by Tymphany) at a distance of 15 cm. The output of the speaker was measured with a free-field microphone (4939-A-011, Brüel & Kjӕr) placed at the same location as the mouse’s head facing the speaker. The output was read in dB sound pressure level (SPL). The pure tones were arranged from 0.5 to 42 kHz in 1/12 octave steps. The duration was 100 ms including a 6 ms cosine^2^ onset/offset ramp and an inter-tone interval of 1300 ms. To align the acoustic stimuli timeline with the recording channels, the click channel from the same connection ports on the sound card with the same onset and offset timeline settings as the tone channel was directly connected to an analog-to-digital converter (ADC) port in the Intan USB interface board and the voltage of the acoustic click channel was recorded. Recording sessions lasted up to 2 h. To allow investigation of electrode performance only, the mice were kept under shallow anesthesia, responding with a light withdrawal reflex to the tail or toe pinch but remaining quiet and motionless without indication of pain or distress. Deep anesthesia was maintained with a higher concentration of isoflurane, and the mice showed no response to the tail or toe pinch.

### Data analysis

All electrophysiological data were analyzed using MATLAB (Mathworks) and sorted with Offline Sorter (OFS, Plexon) using the principal components of the waveforms. A 10-20 min dataset with continuous recording was used for analysis unless otherwise stated. In brief, the raw recording data was filtered using Butterworth bandpass filters (filtfilt function in MATLAB) at 0.1-300 Hz for the local field potential (LFP) and 300-9000 Hz for action potential (AP) extraction. A custom MATLAB script was used to create the power density spectrum in the frequency domain and the spectrogram in time domain based on the raw data to check the recording quality and noise. The amplitude of action potential was calculated using peak-to-peak detection. The signal-to-noise ratio (SNR) was expressed in decibels from 20 × log (RMS(S)/RMS(N)), where S is the waveform peak and N is the noise of the recording after removing the detected spikes. The SNR was evaluated based on the average spike amplitude over a 10-min recording. For sorting and clustering into OFS, the raw recording data was converted to a binary data format and imported to OFS. The data was then filtered with a 2-pole Butterworth high-pass filter at a cutoff frequency of 500 Hz. Spikes at different channels were detected and extracted in a 1.13 ms time window with a prethreshold period of 0.26 ms and dead time of 1.03 ms. All the spike sorting results were triangulated on four neighboring channels. The 16-channel NeuroEdge was grouped in a 4×4 array, and timestamp comparison was applied to the original sorted single units to remove duplicated ones. Interspike intervals (ISI), auto-correlograms and cross-correlograms were exported from OFS and plotted using custom MATLAB scripts. The recording data under acoustic stimuli was analyzed using custom MATLAB scripts with the Gaussian spike density (SD) algorithm based on the average neural response in the form of the peristimulus time histogram (PSTH). The response strength to different auditory frequencies was evaluated via the SD peak around the stimulation window compared with the SD during rest time. Allocation of the spikes to the corresponding frequency of the acoustic stimulation was done by comparing the timestamp of spikes in each putative single neuron with the timeline of the trigger channel. Cross-correlation analyses of the simultaneously recorded neurons were conducted using cross-correlograms following standard Pearson correlation coefficient analysis. Spike timestamps of all the neurons were converted into binary format, with ‘1’ indicating firing events and ‘0’ indicating non-firing events, setting the bin size at 1ms. Pairwise spike trains of identified neurons were then plotted into cross-correlograms. Bins within ± 1ms from the time lag zero in cross-correlograms were excluded as they may originate from the same neuron. The interaction strength was calculated as the average bar height of the normalized cross-correlograms in the interaction time window (1-10 ms of positive and negative time lag) relative to the baseline window (50-100 ms of positive and negative time lag). The connectivity matrices of the interaction strength were constructed by computing all pairwise combinations of the identified neurons.

### Immunohistochemistry

To minimize the impact of probe size on immunohistochemistry (IHC) results, mice were implanted with NeuroEdge and NeuroEdge with thin coating of 100 nm Au/1-2 µm PEDOT:PSS for fair comparison. After three months of implantation, the mice were perfused with saline and 4% paraformaldehyde (PFA) (Sigma-Aldrich) in 0.1 M PB. The fixed brain was extracted and post-fixed in 4% PFA overnight at 4 °C, then immersed in 30% sucrose in 0.1 M PB until the brain sunk to the bottom. The brain was frozen with OCT and sectioned to 45 µm lateral slices at -20 °C using a Cryostat (Leica CM3050s). The individual serial sections were collected into 96- well plates filled with cryoprotection solution and stored at -20 °C until use. The slices were washed with tris-buffered saline (TBS) twice and incubated in a blocking solution containing 0.2% Triton X-100 (Sigma-Aldrich) and 3% donkey serum (GeneTex GTX73205) in TBS for 1 h at room temperature. The slices were then incubated with primary antibodies, including mouse anti-NeuN (MAB377, 1:100, Sigma), chicken anti-GFAP (ab4674, 1:1000, Abcam) and rabbit anti-Iba1 (019-19471, 1:1000, Wako) in blocking solution for one day at 4 °C. The slices were washed with TBS twice for 10 min each, and incubated with secondary antibodies at room temperature for 4 h, including donkey anti-mouse Alexa Fluro 647 (715-605-151, 1:250, Jackson ImmunoResearch), donkey anti-chicken Alexa Fluro 488 (703-545-155, 1:250, Jackson ImmunoResearch) and donkey anti-goat cy3 (705-165-147, 1:250, Jackson ImmunoResearch). After washing with TBS twice, the slices were mounted on polylysine coated glass slide (Epredia Polysine) with Antifade Mounting Medium (Vector Laboratories, H-1200-10) and coverslips. Images were acquired using a confocal laser scanning microscope with a 20× air objective lens (Olympus FV3000). Quantification of the expressed area was analyzed with ImageJ.

## Data availability

All data supporting the findings of this study are available within the article and its Supplementary Information. Additional raw data generated in this work are available from the corresponding author upon reasonable request.

## Code Availability

Offline Sorter, which we used for spike sorting. is available at https://plexon.com/products/offline-sorter/. The remaining analyses were done with custom MATLAB routines, which are available from the corresponding authors upon reasonable request.

## Acknowledgements

The scientific illustrations in Fig. 1 are credited to Z. Goh at the Institute for Health Innovation & Technology, National University of Singapore. Y.L and J.S.H. acknowledge the support from Advanced Research and Technology Innovation Centre (ARTIC) grant (HFM-RP6), the Institute for Health Innovation and Technology. Y.L. acknowledges the support from National University of Singapore Presidential Young Professorship Award (22-4974-A0003). J.S.H. acknowledges the support from the National Research Foundation Singapore (NRFF2017-07), Ministry of Education Singapore (MOE2016-T3-1-004). C.T.L. acknowledges the support from NUS Startup Grant (A-8001301-00-00 and A-0009363-04-00).

## Author contributions

Y.J. and J.S.H. initially conceived the project. Y.J., J.S.H. and Y.L. designed the project. Y.J. fabricated the neural probe, developed the implantation technique, performed probe characterization, in vivo animal surgeries, electrophysiological recording, histology and analyzed neural recording data. Y.J. and J.S.H. designed the figures with Y.L. and Z.X.’s input. Y.J. drew the figures with J.S.H.’s input. C.T., S.A. and Y.J. conducted the MATLAB coding for neural data processing. D.N. and A.T. developed the MATLAB code for generating the sound stimulation recipe. D.N. contributed to correlation analysis. A.T. discussed and supported the auditory experiment and the results analysis. Z.X. contributed to the schematic drawing. R.H. and C.L. contributed to the filtering coding. C.L. contributed to the discussion of neural data processing. J.Y. contributed to coding. C.Y. contributed to frequency analysis coding. S.K. supported the auditory experiment. H.A., Z.L. and B.T. contributed to mechanical measurement. Y.J. prepared the initial manuscript draft. Y.J., J.S.H., Y.L., D.N., Y.K., C.T.L. revised the manuscript. J.S.H., Y.L. and C.T.L. supervised the project. All authors discussed and commented on the manuscript.

## Competing interests

John S. Ho, Yunxia Jin, Yuxin Liu and Chwee Teck Lim are on a patent application filed by the National University of Singapore relating to this work. All the other authors declare no competing interests.

## Supporting Information

Supplementary Information is available for this paper.

## Notes

### Competing Interest Statement

Yunxia Jin, John S. Ho, Chwee Teck Lim and Yuxin Liu are on a patent application filed by the National University of Singapore relating to this work. All the other authors declare no competing interests.

