## Supplementary Information for "Graphene nanoedge electronics for monolithic and chronic recording of local microcircuits at neuronal density"

*Y. Jin et. al.*

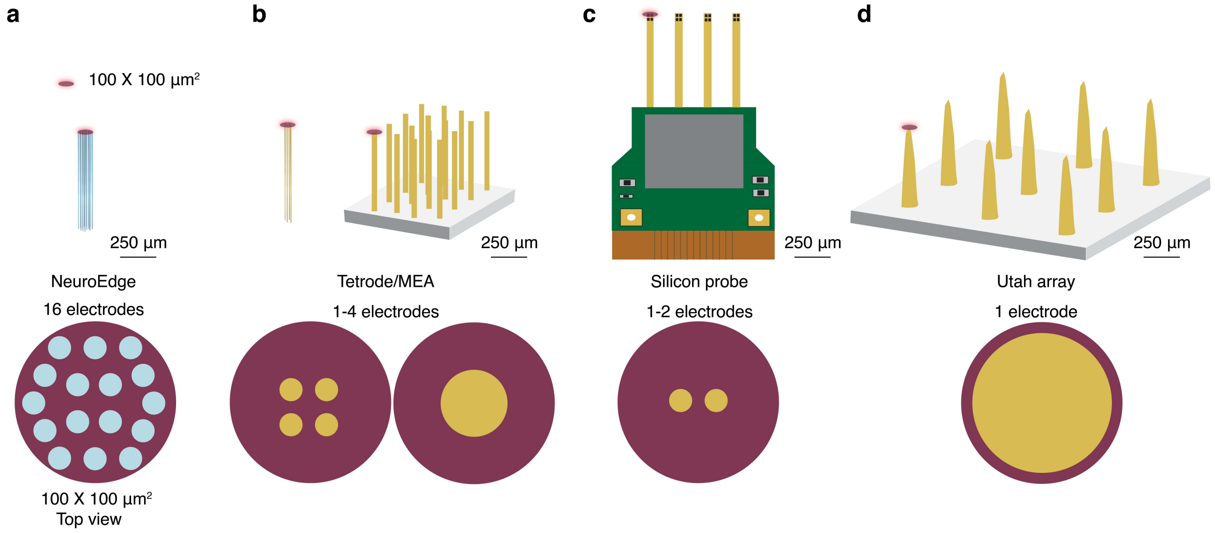


**Fig. S1 Areal density of electrodes. a**, NeuroEdge features 16 electrodes within a compact area of 100 µm × 100 µm. **b**, Microwire arrays or electrode typically include 4 channels from a tetrode metal wire bundle or 1 electrode from single-wire arrays. **c**, Silicon probes typically have 1 or 2 electrodes per 100 µm × 100 µm area. The limitation arises from the 2D planar constraints of semiconductor technology. Typical conductive layers (Au or TiN) of silicon probes have submicron thickness; therefore, their edges are not suitable for neural recording electrodes. **d**, Utah array has 1 electrode per 100 µm × 100 µm area. The interelectrode distance of Utah array is larger than 100 µm.

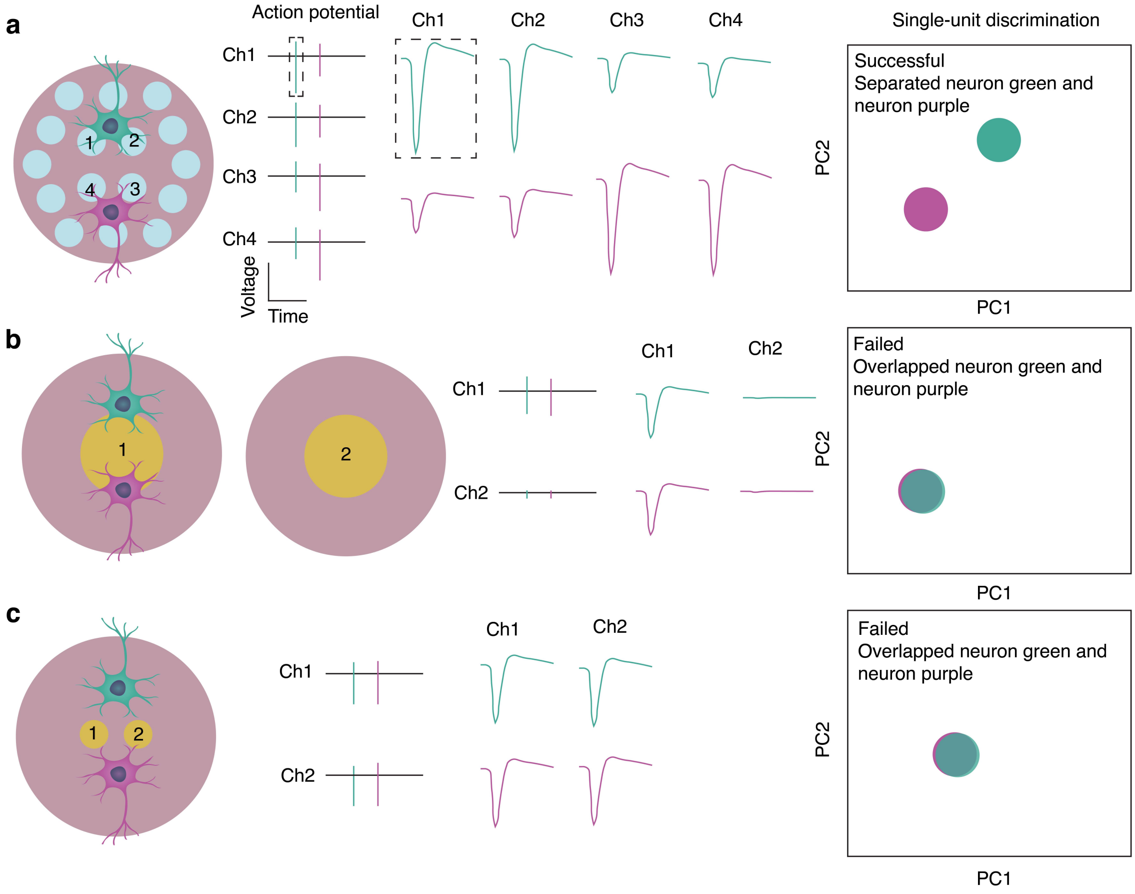


**Fig. S2 Schematics of single-unit spike-sorting across multiple channels and single channel.** **a** NeuroEdge records nearby neurons, e.g., green neuron and purple neurons, on multiple channels with varied spike amplitudes. Both neurons generate similar action potential waveform. The electric field amplitude surrounding a neuron decreases sharply with distance. Thus, the recorded spike amplitudes vary as each channel is positioned at a different distance from the neuron. This provides additional discrimination of the nearby neurons by PCA based spike-sorting, besides merely action potential waveform. **b,** Microelectrode array with large interelectrode spacing records neurons on electrode 1, and no spikes from neuron green and purple on the adjacent electrode 2. If the waveforms of the green neuron and purple neuron are similar, PCA based spike-sorting may fail to sort out the two neurons. **c**, Silicon probe with only 2 electrodes at lateral layer can record near-by neurons on both electrode 1 and electrode 2. But due to the similar positioning to the two electrodes, neuron green and purple may result similar amplitude for both on electrode 1 and 2, and therefore fail to discriminate the two neurons with PCA based spike-sorting.

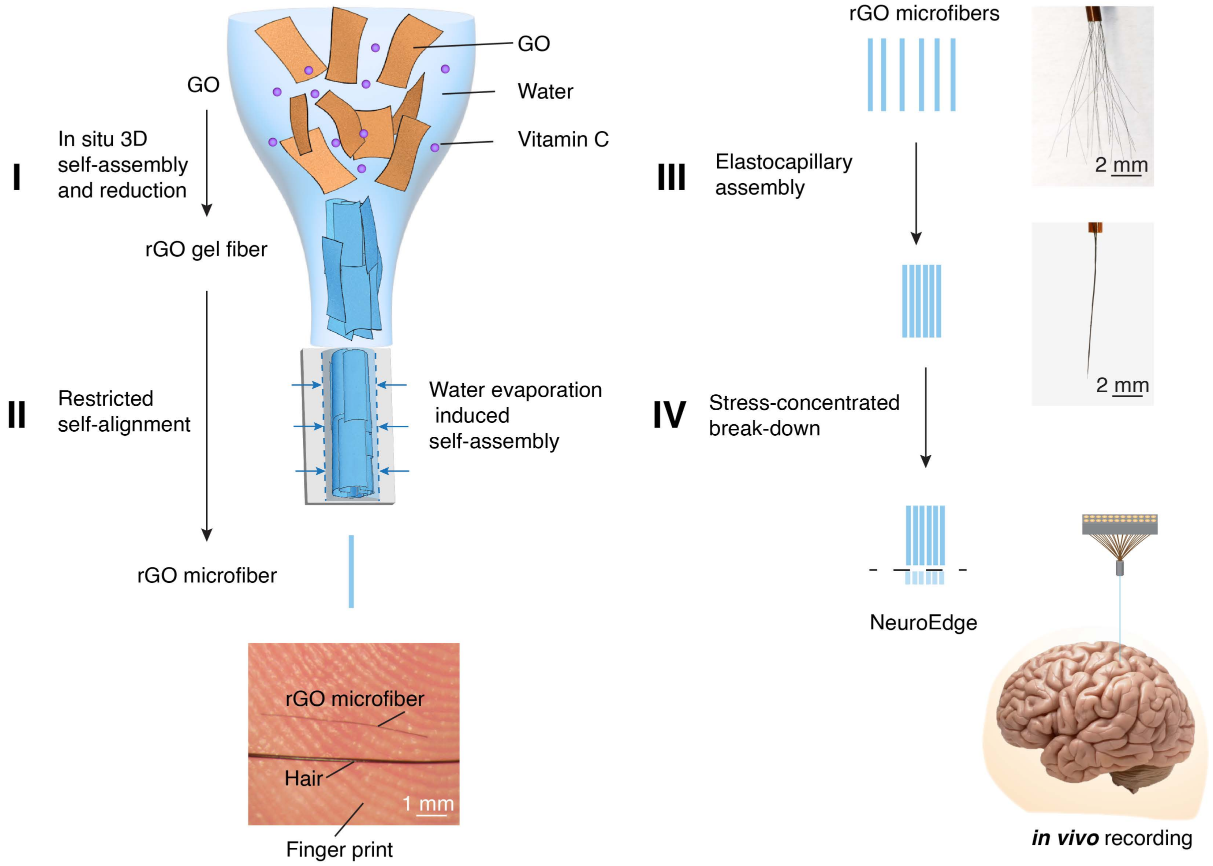


**Fig. S3 Fabrication process of NeuroEdge.** Fabrication includes four key steps, incorporating solution-processable self-assemblies and stress-concentrated break-down. Specifically, in step I, to maximize the in vivo biocompatibility and realize the small form factor while streamline fabrication, solution-processable graphene oxide (GO) sheets were employed to form reduced graphene oxide (rGO) gel fiber through in situ 3D self-assembly and reduction with only water and vitamin C, a natural antioxidant that is widely employed as a food additive and can compete with the highly toxic hydrazine in terms of its reducing capability. Step II involves restricted self-alignment, driven by water evaporation induced self-assembly, to form a straight microelectrode fiber (II) from the gel fiber. The photograph reveals the prepared microfiber is significantly slimmer than the hair and fingerprint. In step III, to maximize array density, elastocapillary self-assembly was employed, merging the parylene C insulated microfibers into a bundle during the withdrawal from a bath of diluted SU-8 solution. The small form factor of microfiber enabled the elastocapillary self-assembly by providing a balanced adhesion energy arising from van der Waals force and probe stiffness. Such assembly cannot be extended to metal wires in similar neuronal size or larger metal probes as their stiffness will exceed the capillary adhesion force, preventing the formation of bundles. The grand challenge of this high-density configuration is to maintain a low impedance while reduce the recording site to neuronal size. Therefore, in step IV we developed a stress-concentrated break down method to expose the bundle tip end. The rGO sheet stack are well exposed preserving the nanoedge, nanotexture and nanotunnels, which accounts for the low impedance. During the whole process, the absence of toxic or organic solvent aligned with small size of probe, prioritizes the biocompatibility essentials for implantable probes. The solution processable technology allows for precise tuning of the probe to any desirable size. The stitched rGO sheets via van der Waals force enable highly flexibility and robustness of the NeuroEdge.

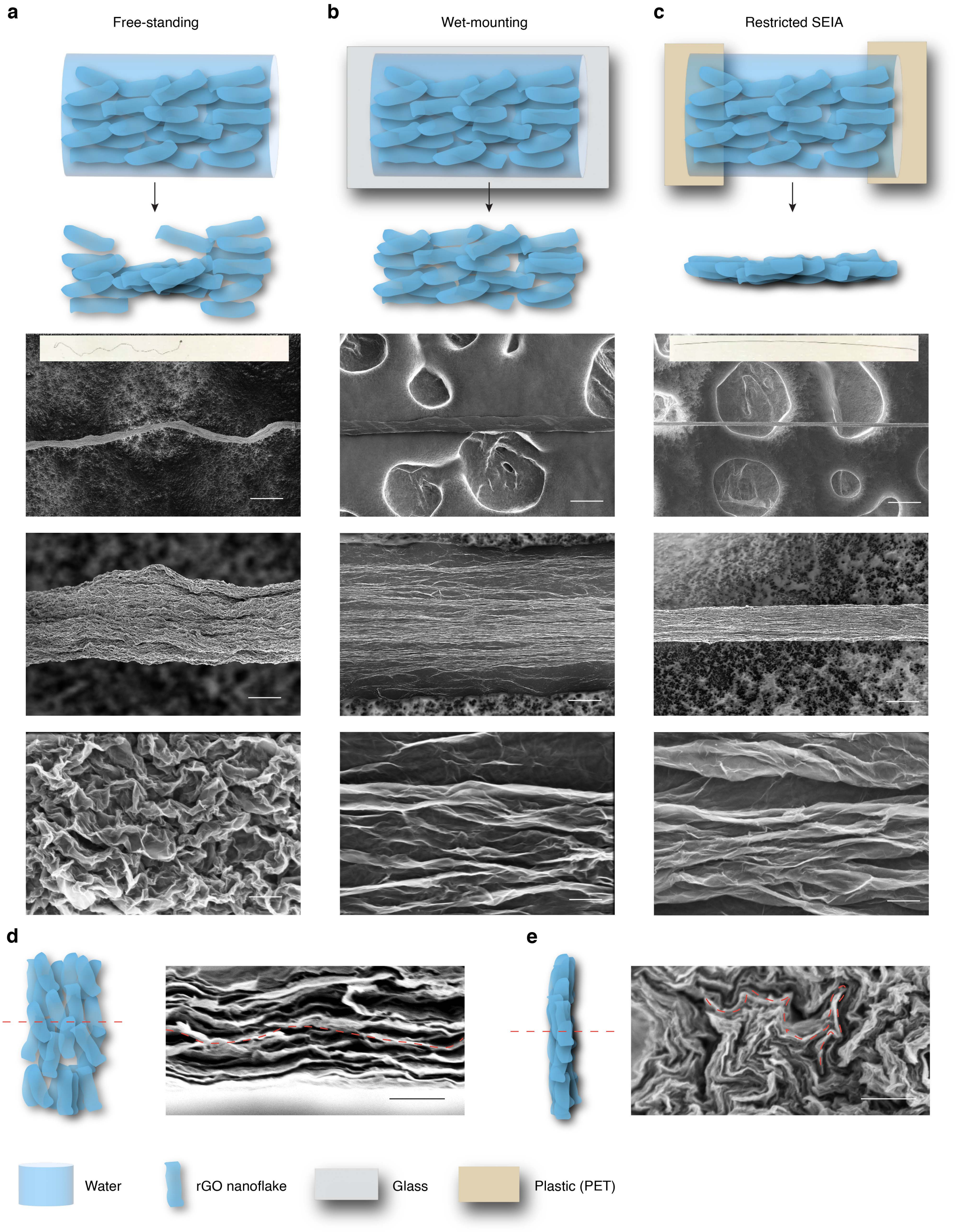


**Fig. S4 Step II of fabrication process of NeuroEdge.** We examined three approaches for creating rGO microfiber from gel fiber with the goal of minimizing axial footprint size and fluctuation in diameter. **a,** Schematic of free-standing approach and the SEM images of the prepared microfiber. The gel fiber was exposed to air and underwent natural evaporation of water within the gel fiber. The evaporation process allows the rGO sheets to contact each other and shrink. However, the uneven evaporation rate, influenced by distinct radii of curvature at different locations along the gel fiber, led to uneven movement of rGO sheets, resulting in varying diameters at different locations. The SEM shows deep folding or crumpling of rGO sheets. **b,** Schematic of wet-mounting approach and the SEM images of the prepared fiber. The gel fiber was placed on a glass substrate. As water evaporated, the high surface tension of water caused the gel fiber to wet the hydrophilic glass, expanding its axial footprint. The bottom layer of rGO sheets were adhered to the glass due to interface tension between water and glass, minimizing crumpling and allowing for the stacking of rGO sheets layer-by-layer. The SEM images show the outer rGO sheets appear much flatter compared to those in a. with a footprint of 70 µm. **c,** Schematic of restricted solvent evaporation induced self-assembly (SEIA) and the SEM images of the resulting fiber. The gel fiber was anchored at both ends on a plastic substrate, and underwent solvent evaporation induced self-assembly of the rGO sheets along the anchoring direction. The absence of interface tension allowed the rGO sheets to freely assembly while being confined to the mounting direction, resulting in a small diameter and a straight longitudinal alignment. The SEM images showed the diameter is only 12 µm which is 5-fold smaller compared to b. **d-e,** Schematics of viewing direction and the SEM images from cross-section for fiber that was prepared by methods in b (**d**) and c (**e**). The folding of rGO sheets in e from restricted SEIA method resulted in a wavy pattern, while flattened rGO sheets from the wet-mounting method showed a layered pattern. It highlights the advantage of the method in c for reducing the diameter. Scale bar is 100, 10 and 0.5 µm for SEM images from top to bottom in a, b and c, and 0.5 µm in d and e.

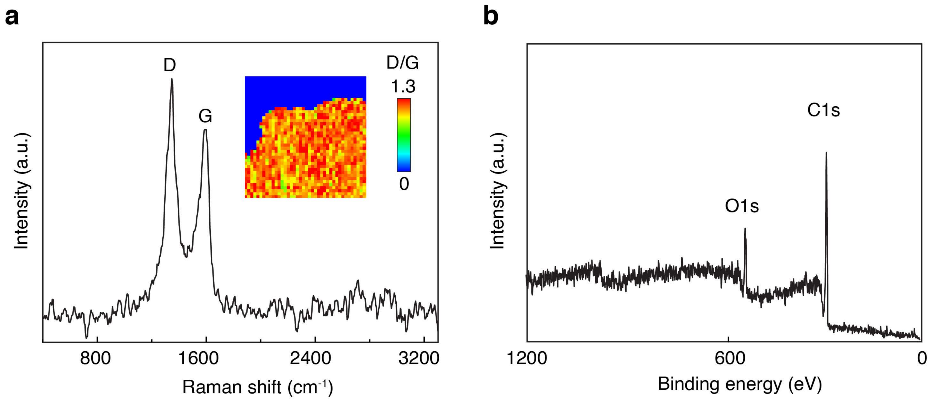


**Fig. S5 a-b,** Raman characterization (**a**) and X-ray photoelectron spectroscopy (XPS) (**b**) for NeuroEdge. The inset in a is the Raman scan of the exposed tip end of NeuroEdge. The D/G ratio is predominantly around 1.3. The XPS result showed a significantly increased ratio of carbon-to-oxygen compared to graphene oxide (GO). It indicates that the GO has been successfully converted to reduced graphene oxide (rGO).

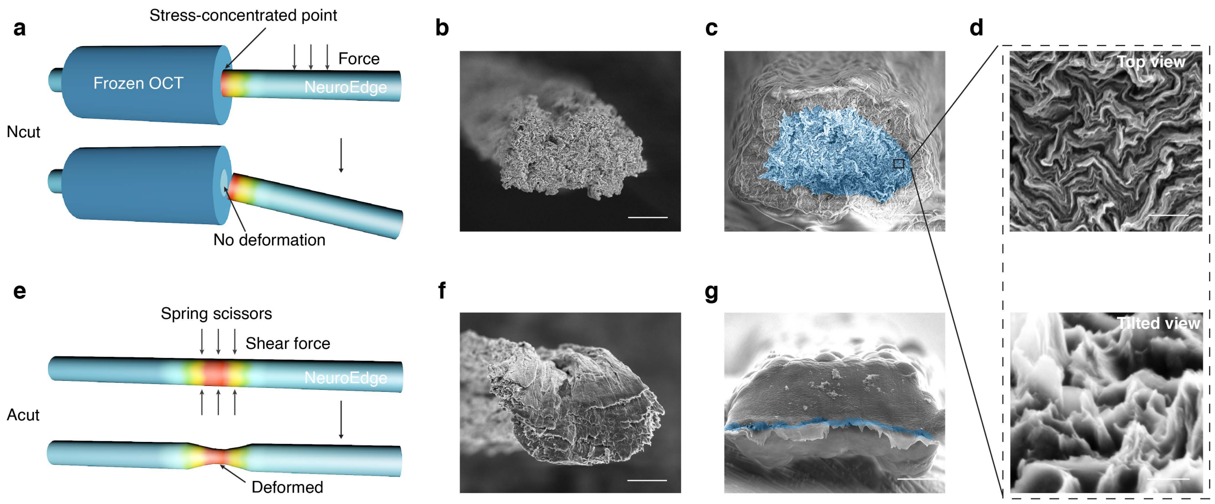


**Fig. S6 Stress-concentrated breaking-down technique to expose the electrode tip. a,** Schematics of stress-concentrated breaking-down method (Natural cut (Ncut)). It involves partially coating the frozen optimal cutting temperature (OCT) compound on probe, increasing the diameter at one end, and hence causing significant bending stiffness difference between coating and uncoating part. It leads to naturally break-down of the probe from the coating-uncoating point upon applying force, leaving a sharp exposure of graphene edge with well-preserved nanotexture and nanotunnels. **b-c,** SEM images of cross-section of the bare microfiber (**b**) and microfiber with parylene C (**c**) exposed by Ncut. **d,** Enlarged SEM images in c by viewing from top and tilted angle. It shows wavy nanotunnels and nanoedges. **e,** Schematics of scissors cutting in air (Acut) directly. It causes the deformation of the microfiber due to shear force from the scissors cutting. **f-g,** SEM images of cross-section of bare microfiber (**f**) and microfiber with parylene C (**g**) that were exposed by Acut. It shows deformed microfiber and the parylene C which reduced the exposed electrode area resulting in impedance spiking. Scale bar is 5 µm in b, c, f and g, 500 nm on the top and 250 nm at bottom in d.

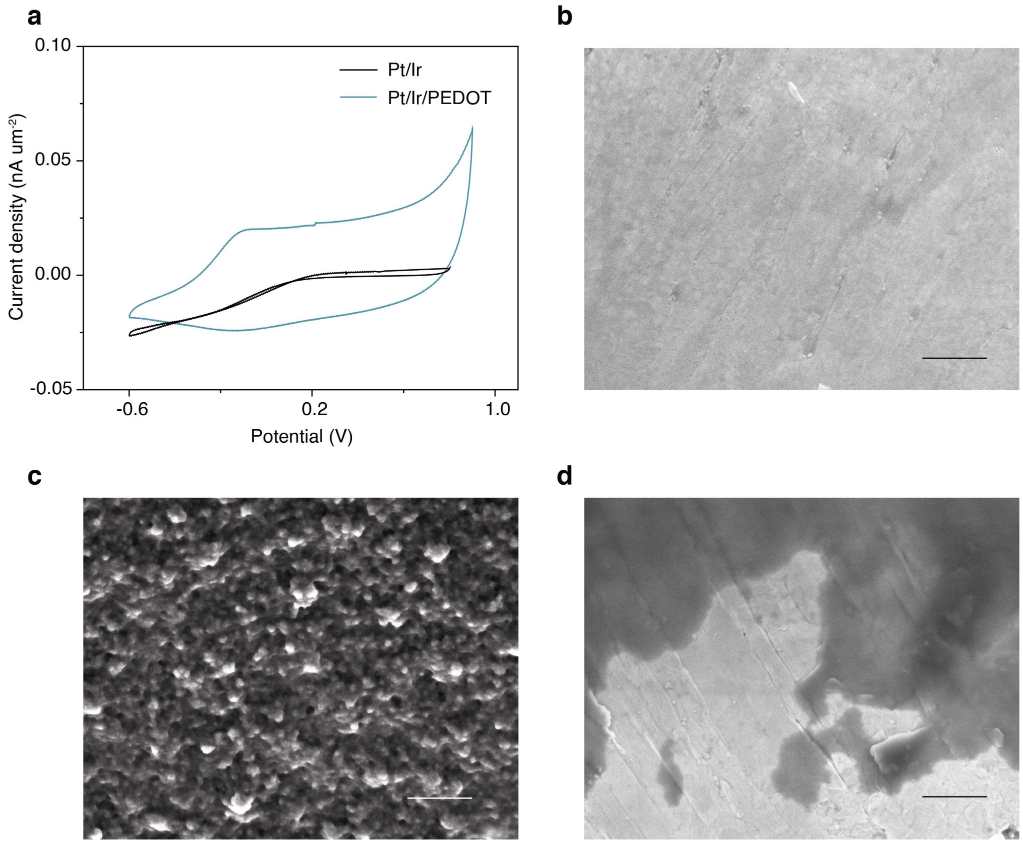


**Fig. S7 Poly(3,4-ethylenedioxythiophene) polystyrene sulfonate (PEDOT:PSS) plating on Pt/Ir metallic electrode**. **a**, Cyclic voltammetry (CV) characteristics of bare Pt/Ir and PEDOT:PSS modified Pt/Ir. **b-d**, SEM images of bare Pt/Ir (**b**), Pt/Ir with deposition of PEDOT:PSS before (**c**) and after electrochemical stress (current pulses). It indicates the successful deposition of PEDOT:PSS on the surface of Pt/Ir electrode, and the detachment of the PEDOT:PSS after electrochemical stress (current pulses). Scale bar is 500 nm.

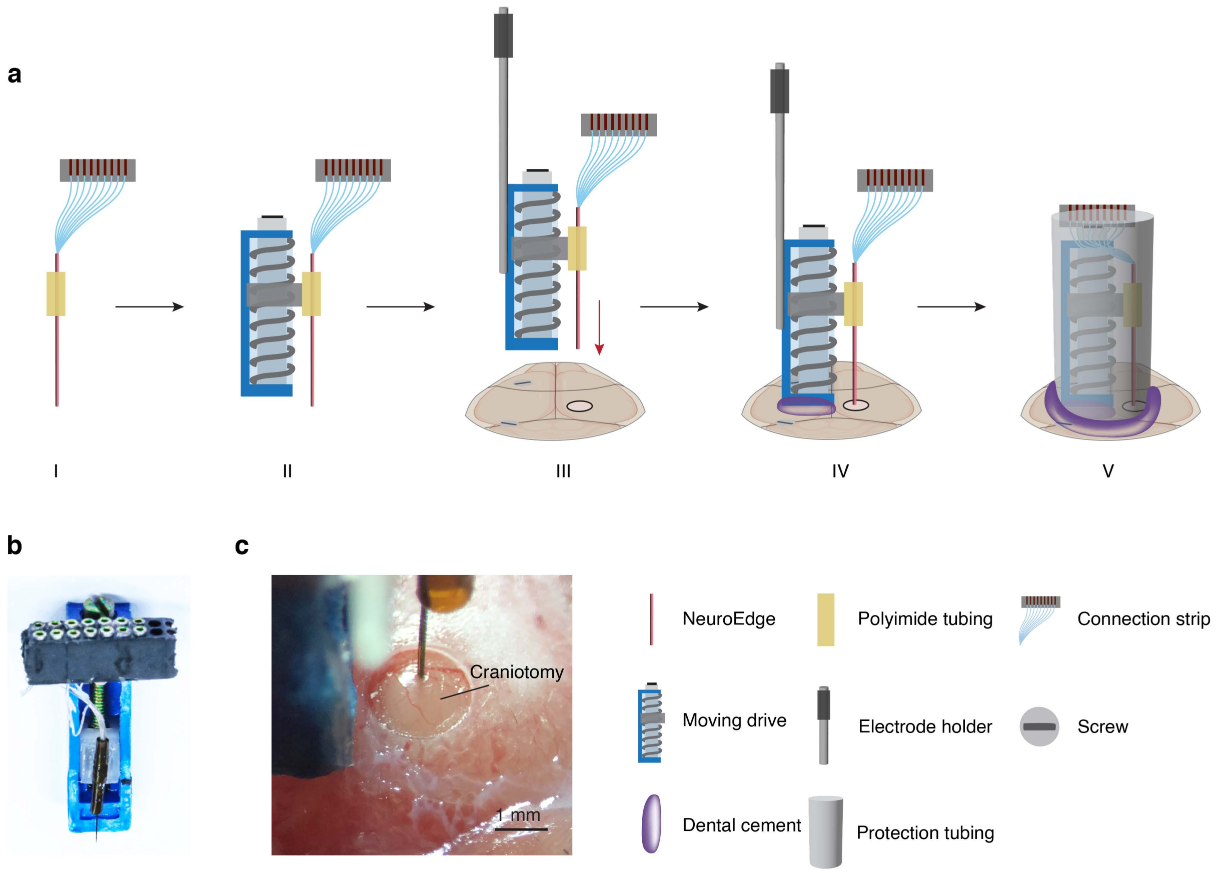


**Fig. S8 Implantation of NeuroEdge**. **a,** Schematics of fabrication and implantation of the NeuroEdge drive into mouse brain. **b,** Photograph of one representative probe. **c,** Photograph of the implantation of the probe into brain. NeuroEdge was attached onto a movable drive. With a clockwise turn of the screw on drive, the electrode was loaded down around 180 μm.

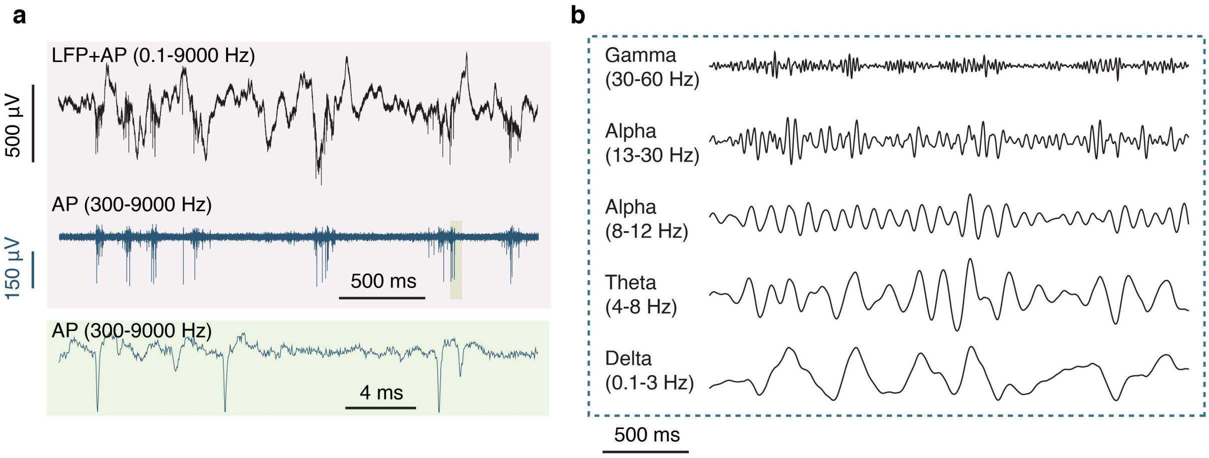


**Fig. S9 Electrophysiological signal recorded by NeuroEdge. a**, Representative local field potential (LFP), action potential (AP). **b,** Representative LFPs at different frequencies.

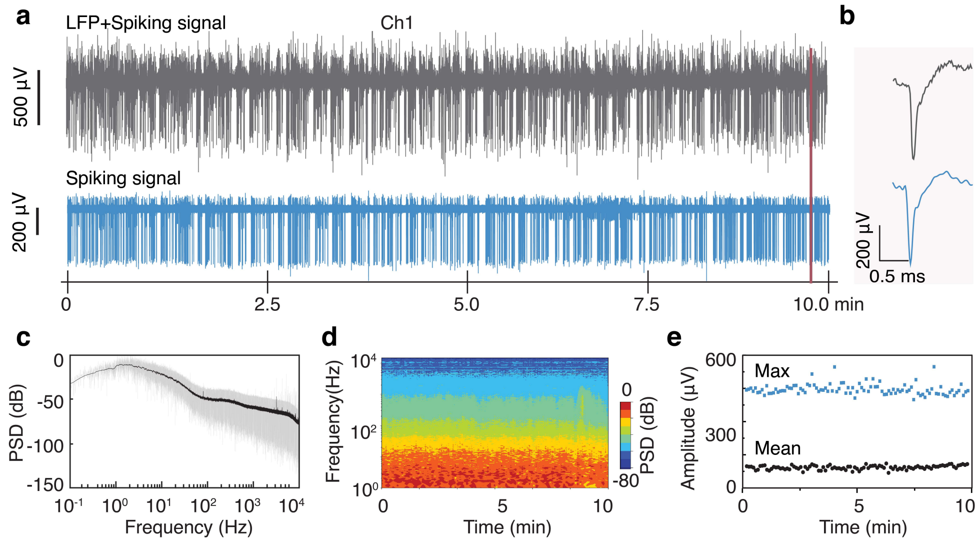


**Fig. S10. Neural recording from NeuroEdge**. **a**, Representative 10-min neural recording from channel 1 (Ch1) in the cortex shows both local field potential (LFP) (0.1-9000 Hz) and spontaneous spiking signal (300-9000 Hz). **b**, A typical waveform before and after high-pass (HP) filtering at 300 Hz in **a** marked by a red line, displaying clear depolarization, repolarization and relative refractory periods. **c, d**, Power spectrum density (PSD) (**c**), and spectrogram (**d**) of the recorded signal (0.1-9000 Hz). **e,** Amplitude in maximum and average of the recorded signal after 300-9000 Hz band-pass filtering.

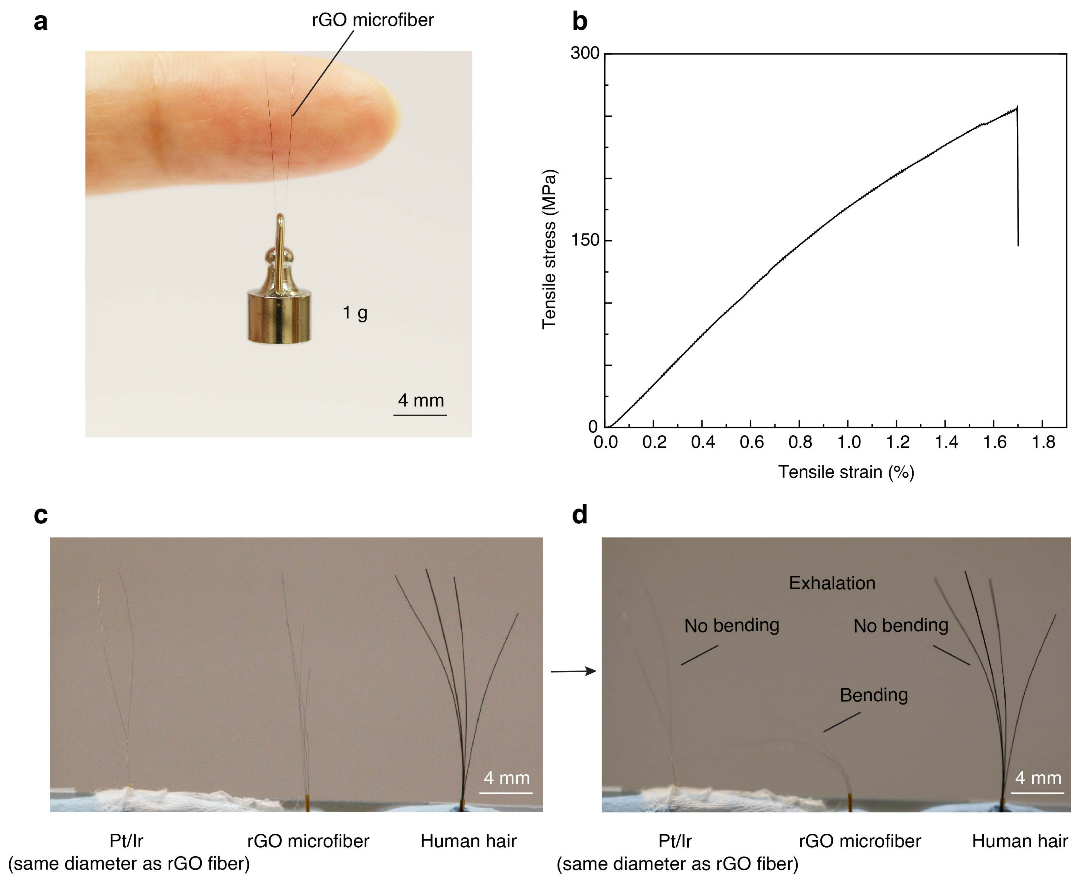


**Fig. S11 Mechanical property of rGO microfiber. a**, Photograph of rGO fiber supporting a weigh of 1 g. **b,** Tensile test of the microfiber. **c-d**, Platinum/iridium10% (Pt/Ir) wire, rGO microfiber, hair before (c) and after (d) subject to airflow with the same intensity. Pt/Ir wire and hair remain still without bending, while rGO microfiber exhibits bending under airflow.

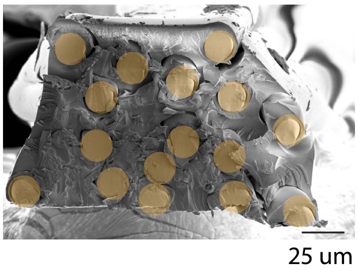


**Fig S12. SEM image of 17.5 um Pt/Ir metal bundle**. It was fabricated using a similar process as NeuroEdge, but has over twice probe area in the cross-sectional direction. This is due to reduced conformal deformation during the self-assembly process, resulting from its higher modulus.

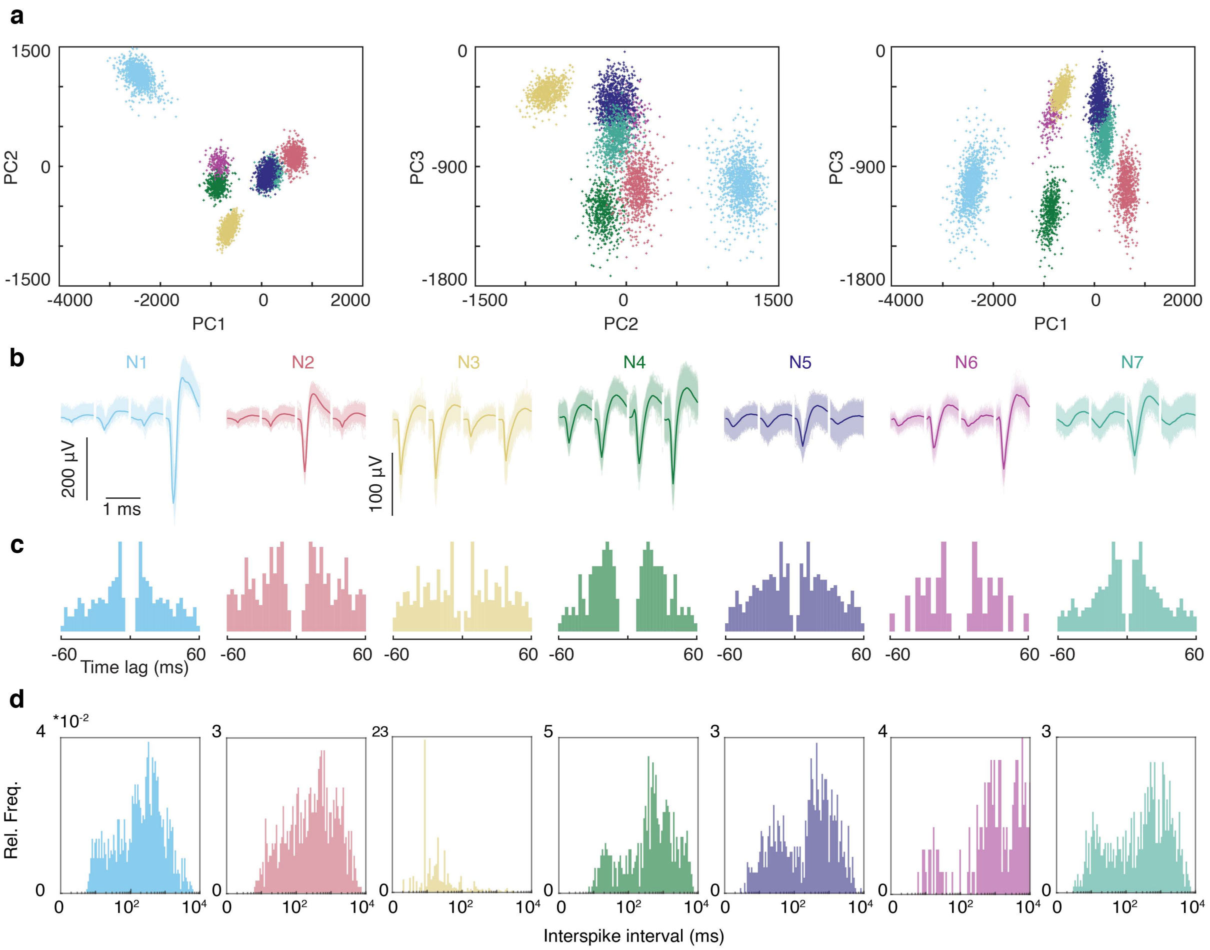


**Fig. S13 Spike sorting of the 4-channel NeuroEdge. a,** PCA projections of the single units. **b,** The waveforms of the single units. The solid lines represent the average waveform. **c**, Auto-correlogram of each unit. **d**, Interspike internal (ISI) of each unit.

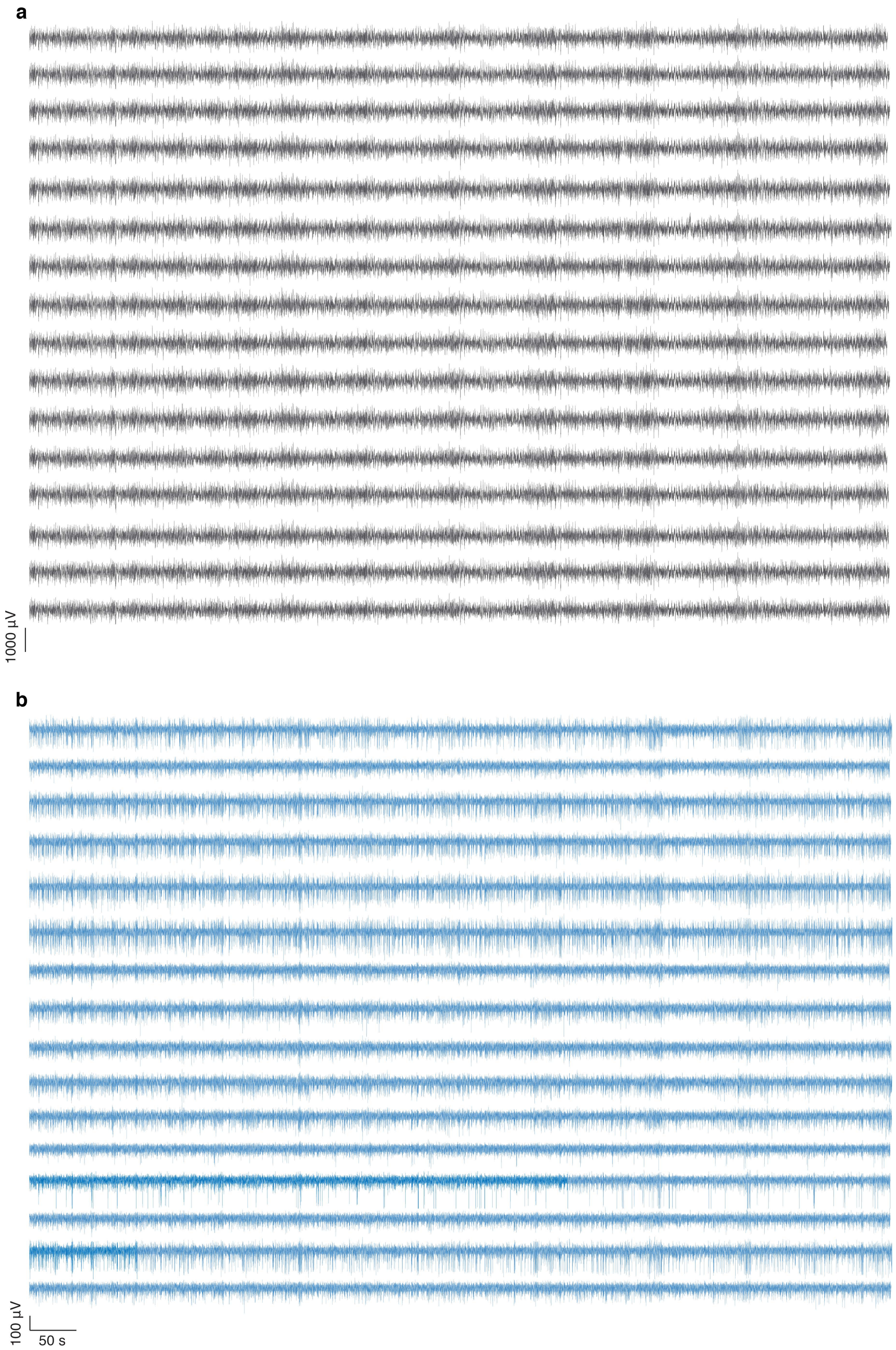


**Fig. S14 Electrophysiological signal recorded by 16-channel NeuroEdge**. **a,** LFP and AP, **b,** AP.

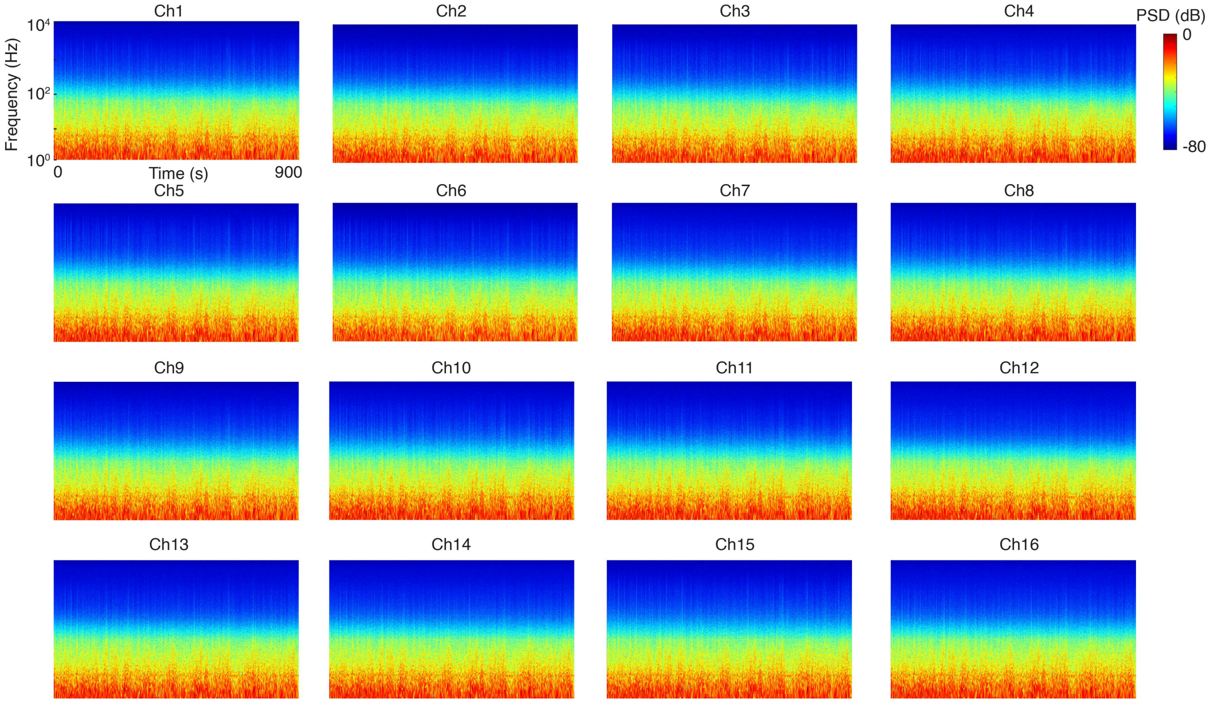


**Fig.S15** Power spectrum density (PSD) of the recorded signal including LFP and AP on each channel.

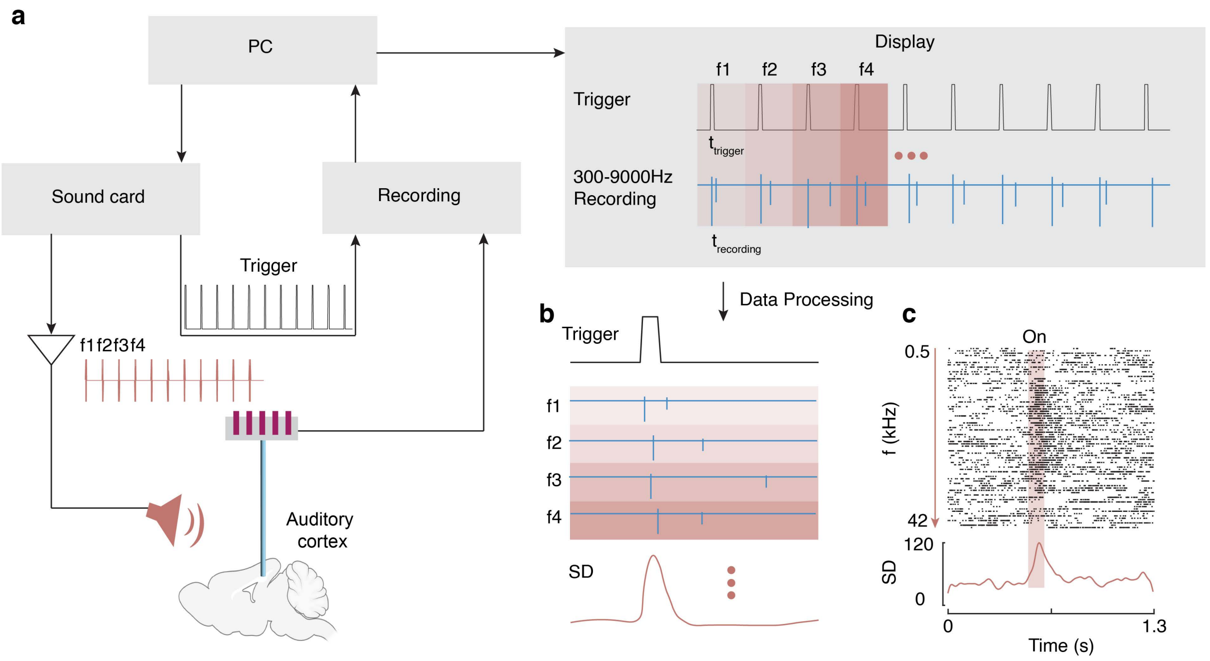


**Fig. S16 Auditory stimulation setup and data processing. a,** Schematics of the auditory stimulation setup. It contains two channels, with the trigger channel marking the timestamp of the stimulation tone aligned with the recorded signal. Each frequency is stimulated for 100 ms, and interstimulus interval is 1.3 s to leave enough rest time for the neuron to recover from the last stimulus. **b,** Schematics of data processing. The filter signal was realigned by interval of 1.3 s, and each corresponds to a sound frequency. **c,** Peristimulus raster plot.

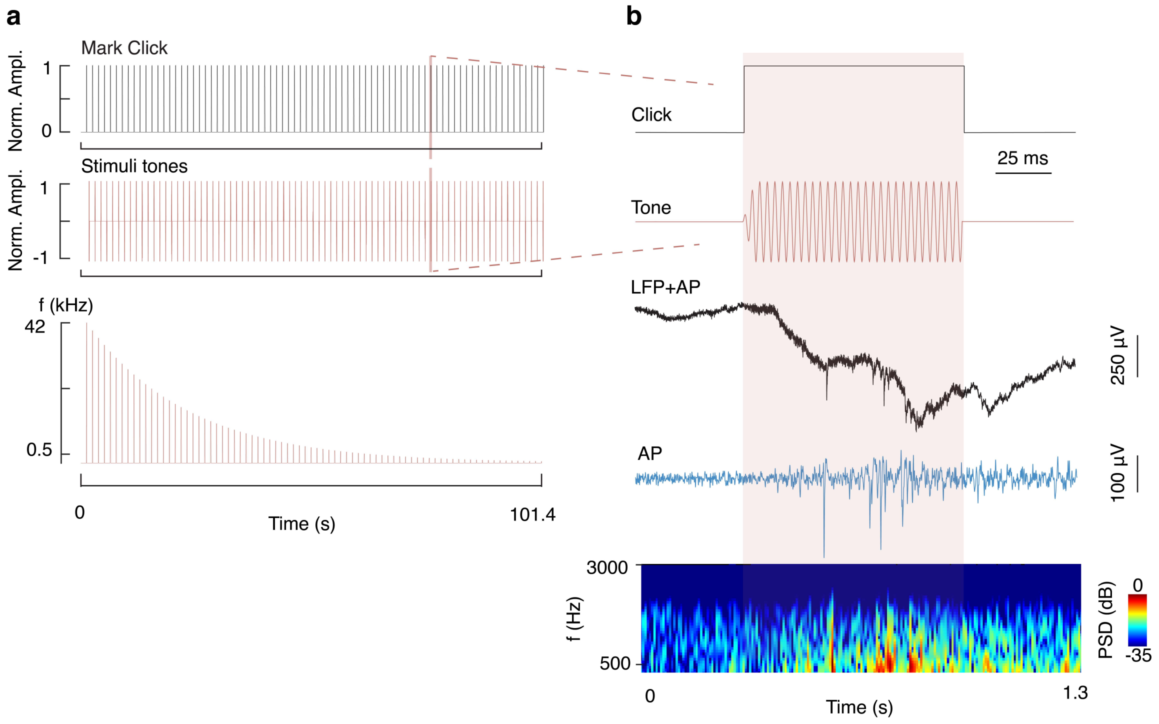


**Fig. S17** Auditory stimulation response. **a**, Auditory stimulation recipes. **b**, Representative neural recording and its spectrogram during the stimulus duration in **a** marked by a red background.

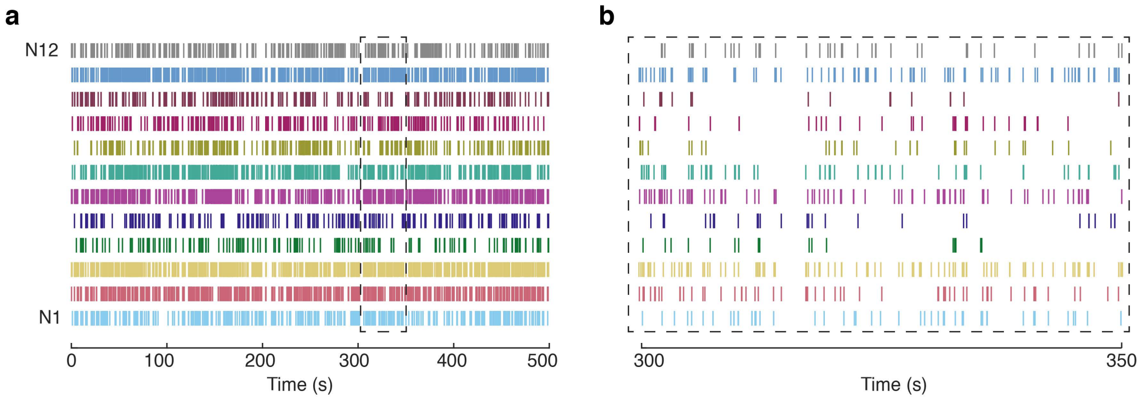


**Fig. S18 a,** Spiking raster from each single unit under acoustic stimulation. **b,** Enlarged view of the spiking raster marked within the dashed rectangular area in a.

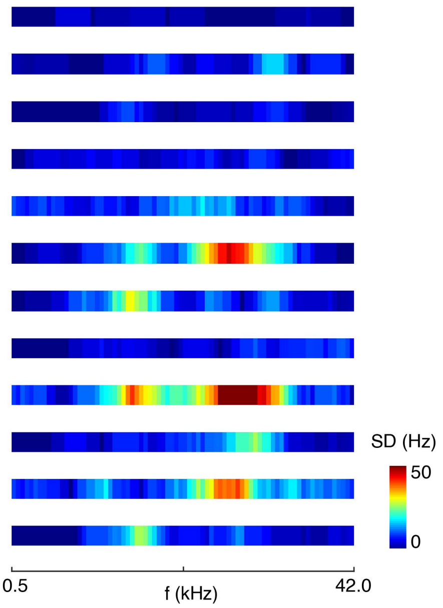


**Fig. S19** Frequency response distribution of the a representative single unit.

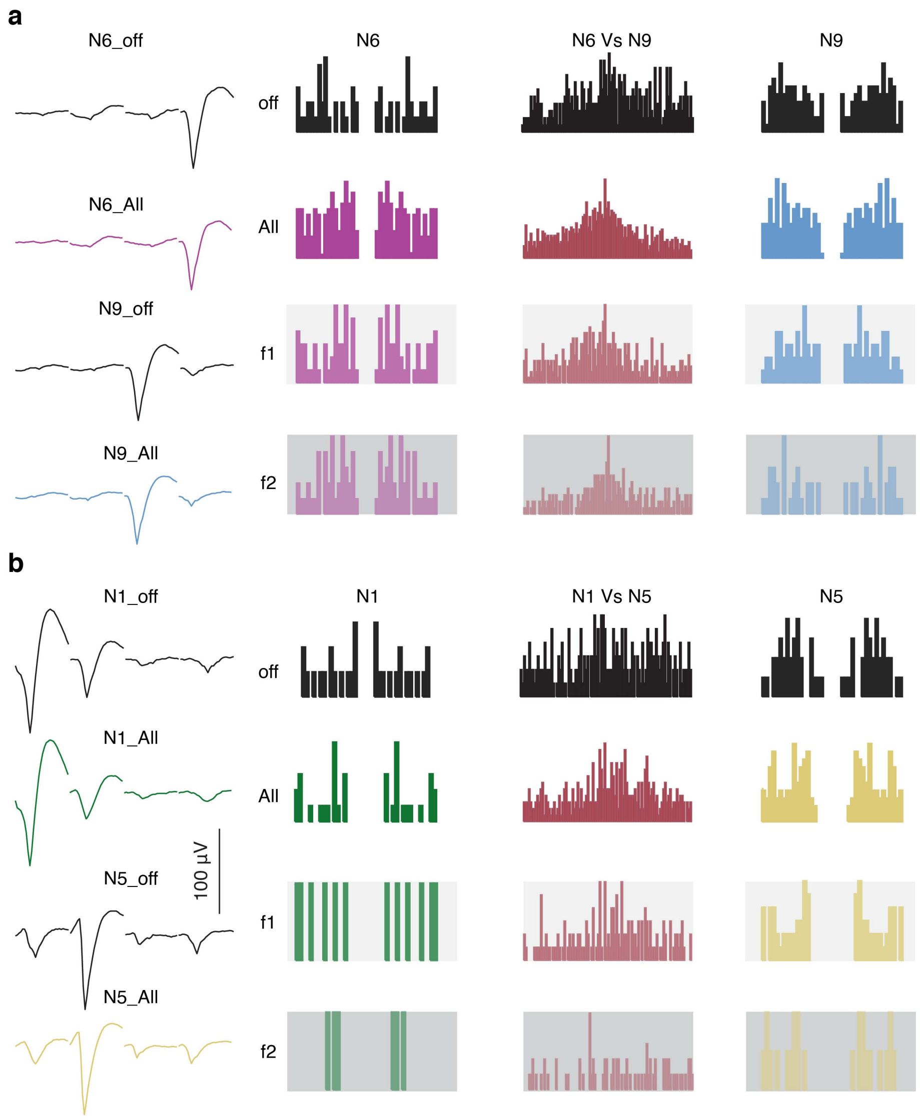


**Fig. S20 Functional connectivity between neurons. a,** left are average waveforms of the neuron 6 (N6) and neuron 9 (N9) without and with acoustic stimulation. Two neurons show distinct waveforms across different channels, confirming they are well-isolated neurons. The waveforms remain consistent for the same neuron before and after acoustic stimulation, indicating accurate tracking of the same individual neuron with NeuroEdge. Right are auto- and cross-correlograms of the neurons under absent stimuli and stimuli at all frequencies, lower frequency f1 and higher frequency f2. **b,** single unit waveform, auto- and cross-correlograms for N1 versus N5.
